# Maelstrom Represses Canonical Polymerase II Transcription within Bi-Directional piRNA Clusters in *Drosophila melanogaster*

**DOI:** 10.1101/344572

**Authors:** Timothy H. Chang, Eugenio Mattei, Ildar Gainetdinov, Zhiping Weng, Phillip D. Zamore

## Abstract

In *Drosophila*, 23–30 nt long PIWI-interacting RNAs (piRNAs) direct the protein Piwi to silence germline transposon transcription. Most germline piRNAs derive from dual-strand piRNA clusters, heterochromatic transposon graveyards that are transcribed from both genomic strands. These piRNA sources are marked by the Heterochromatin Protein 1 homolog, Rhino (Rhi), which facilitates their promoter-independent transcription, suppresses splicing, and inhibits transcriptional termination. Here, we report that the protein Maelstrom (Mael) represses canonical, promoter-dependent transcription in dual-strand clusters, allowing Rhi to initiate piRNA precursor transcription. In addition to Mael, the piRNA biogenesis factors Armitage and Piwi, but not Rhi, are required to repress canonical transcription in dual-strand clusters. We propose that Armitage, Piwi, and Mael collaborate to repress potentially dangerous transcription of individual transposon mRNAs within clusters, while Rhi allows non-canonical transcription of the clusters into piRNA precursors without generating transposase-encoding mRNAs.

## INTRODUCTION

In *Drosophila melanogaster*, 23–30 nt long PIWI-interacting RNAs (piRNAs) direct transposon silencing by serving as guides for Argonaute3 (Ago3), Aubergine (Aub), and Piwi, the three fly PIWI proteins (Aravin et al., 2001; Girard et al., 2006; Aravin et al., 2006; Grivna et al., 2006; Lau et al., 2006; Vagin et al., 2006). In the germ cell cytoplasm, Aub and Ago3 increase the abundance of their guide piRNAs via the ping-pong cycle, an amplification loop in which cycles of piRNA-directed cleavage of sense and antisense transposon-derived long RNAs generate new copies of the original piRNAs—so-called secondary piRNAs—in response to transposon transcription (Brennecke et al., 2007; Gunawardane et al., 2007). In addition to amplifying piRNAs, the ping-pong pathway also produces long 5′ monophosphorylated RNA that enters the primary piRNA pathway, generating head-to-tail strings of piRNAs bound to Piwi, and, to a lesser extent, Aub (Han et al., 2015a; Mohn et al., 2015; Senti et al., 2015; Wang et al., 2015). Unlike Ago3 and Aub, Piwi acts in both the germline and the adjacent somatic follicle cells to represses transposon transcription rather than to cleave their transcripts (Cox et al., 2000; Brennecke et al., 2007; Malone et al., 2009; Klenov et al., 2011). Nuclear Piwi is believed to bind nascent RNA transcripts, and, by binding the protein Panoramix, tethers the histone methyltransferase SETDB1 to transposon-containing loci. SETDB1, in turn, trimethylates histone H3 on lysine 9 (H3K9me3), a modification required to create repressive constitutive heterochromatin (H3K9; Rangan et al., 2011; Sienski et al., 2012; Donertas et al., 2013; Le Thomas et al., 2013; Muerdter et al., 2013; Ohtani et al., 2013; Rozhkov et al., 2013; Klenov et al., 2014; Sienski et al., 2015; Yu et al., 2015b; Aravin et al., 2008; Shpiz et al., 2011; Post et al., 2014).

piRNA precursor RNAs are transcribed from piRNA clusters, heterochromatic loci that comprise transposons and transposon fragments that record a species’ evolutionary history of transposon invasion (Brennecke et al., 2007; Lagarrigue et al., 2013; Aravin et al., 2008; Fu et al., 2018b). *Drosophila* piRNA clusters can be uni-strand, transcribed from one genomic strand, or dual-strand, transcribed from both genomic strands. Uni-strand clusters, such as the ∼180 kbp *flamenco* (*flam*) locus, silence transposons in somatic follicle cells (Pelisson et al., 1994; Prud’homme et al., 1995; Robert et al., 2001; Sarot et al., 2004; Mevel-Ninio et al., 2007; Pelisson et al., 2007), whereas dual-strand clusters, such as the ∼250 kbp *42AB* locus, predominate in the germline (Malone et al., 2009). Some uni-strand clusters, such as *cluster2*, are active in both tissues.

Standard, promoter-initiated, RNA polymerase II (Pol II) transcription generates spliced, polyadenylated precursor piRNAs from *flam* (Robert et al., 2001; Mevel-Ninio et al., 2007; Goriaux et al., 2014). In contrast, dual-strand clusters generally lack conserved promoters. Instead, the Heterochromatin Protein 1 homolog Rhino (Rhi) binds to the H3K9me3 present on the piRNA clusters, to which it can tether additional proteins (Lachner et al., 2001; Cogoni and Macino, 1999; Klattenhoff et al., 2009; Zhang et al., 2012; Le Thomas et al., 2014; Mohn et al., 2014; Zhang et al., 2014; Yu et al., 2015a). One Rhi-associated protein, Moonshiner (Moon), is a germline-specific TFIIA-L paralog that allows Pol II to initiate transcription without promoter sequences, allowing every bound Rhi to be a site of potential transcriptional initiation (Andersen et al., 2017). Another Rhi-binding protein, Cutoff (Cuff), suppresses splicing and transcriptional termination (Pane et al., 2011; Mohn et al., 2014; Zhang et al., 2014; Chen et al., 2016). Thus, Rhi promotes “incoherent” transcription, RNA synthesis initiating at many sites throughout both strands of a dual-strand cluster, in contrast to the “coherent,” promoter-dependent transcription of *flam* and conventional protein-coding genes.

Maelstrom (Mael), a protein with HMG- (Findley, 2003) and MAEL- (Zhang et al., 2008a) domains, has been suggested to play multiple roles in *Drosophila* oogenesis and mouse spermatogenesis, including repression of the microRNA miR-7, transposon silencing, heterochromatin formation, and piRNA production. Here, we report that Mael suppresses coherent transcription within dual-strand piRNA clusters. In *mael* mutant ovaries, piRNA cluster heterochromatin organization and transcription are largely unaltered. However, without Mael, Piwi, or Armitage (Armi), a core piRNA biogenesis protein, transcription initiates from canonical Pol II promoters within dual-strand clusters such as *42AB*. Although Rhi and Cuff are required for incoherent transcription of dual-strand piRNA clusters, they are dispensable for repression of canonical transcription in the clusters. We propose that Mael, Armi, Piwi, and piRNAs collaborate to repress canonical dual-strand cluster transcription, while Rhi serves both to create a transcriptionally permissive chromatin environment and to support incoherent transcription of both DNA strands of dual-strand clusters. Thus, Mael represses promoter-driven transcription of individual, potentially active, transposons embedded within dual strand-clusters, allowing Rhi to transcribe such transposon sequences into intron-containing piRNA precursors with little potential to be translated into transposon-encoded proteins required for transposition.

## RESULTS

### Maelstrom Represses Transcription in Dual-Strand Clusters

Without Mael, both somatic and germline transposons produce long RNA transcripts (Sienski et al., 2012; Muerdter et al., 2013; Pek et al., 2009)) while protein-coding genes are largely unaffected (Figure S1). In the germline of *mael^M391/r20^* ovaries, steady-state RNA abundance from telomeric transposons increased >360-fold for *HeT-A* and *TART* and ∼49-fold for *TAHRE* (*n* = 3; Figure S1). Intriguingly, RNA increased >13-fold from two individual *gypsy12* long terminal repeat (LTR) transposon insertions: one in the dual-strand piRNA cluster *42AB* (at 42A14; hereafter *gypsy12^42AB^*) and one in the dual-strand piRNA cluster *cluster62* (at 40F7; hereafter *gypsy12^cluster62^*; Figures 1 and S1). The same two *gypsy12* elements are also desilenced in Rhi- or Cuff-deficient ovaries (Zhang et al., 2014; Mohn et al., 2014). As in *rhi* and *cuff*, but unlike wild-type, RNA from the two *gypsy12* LTRs was spliced in *mael* mutant ovaries. The increase in steady-state *gypsy12* LTR RNA at these two loci in *mael^M391/r20^* mutants reflects a concomitant increase in nascent transcription as measured by global run-on sequencing (GRO-seq; Figure 1; Core et al., 2008). Lysine 4 trimethylation of histone H3 (H3K4me3), a chromatin mark associated with active, promoter-driven transcription (Bernstein et al., 2002; Santos-Rosa et al., 2002; Schneider et al., 2004), also increased at both the *gypsy12* element in *42AB* and in *cluster62* (>3-fold and >9-fold, respectively, *n* = 2; Figure 1). These data suggest that in the absence of Mael, RNA polymerase II initiates canonical transcription within the *gypsy12* LTR.

**Figure 1.**
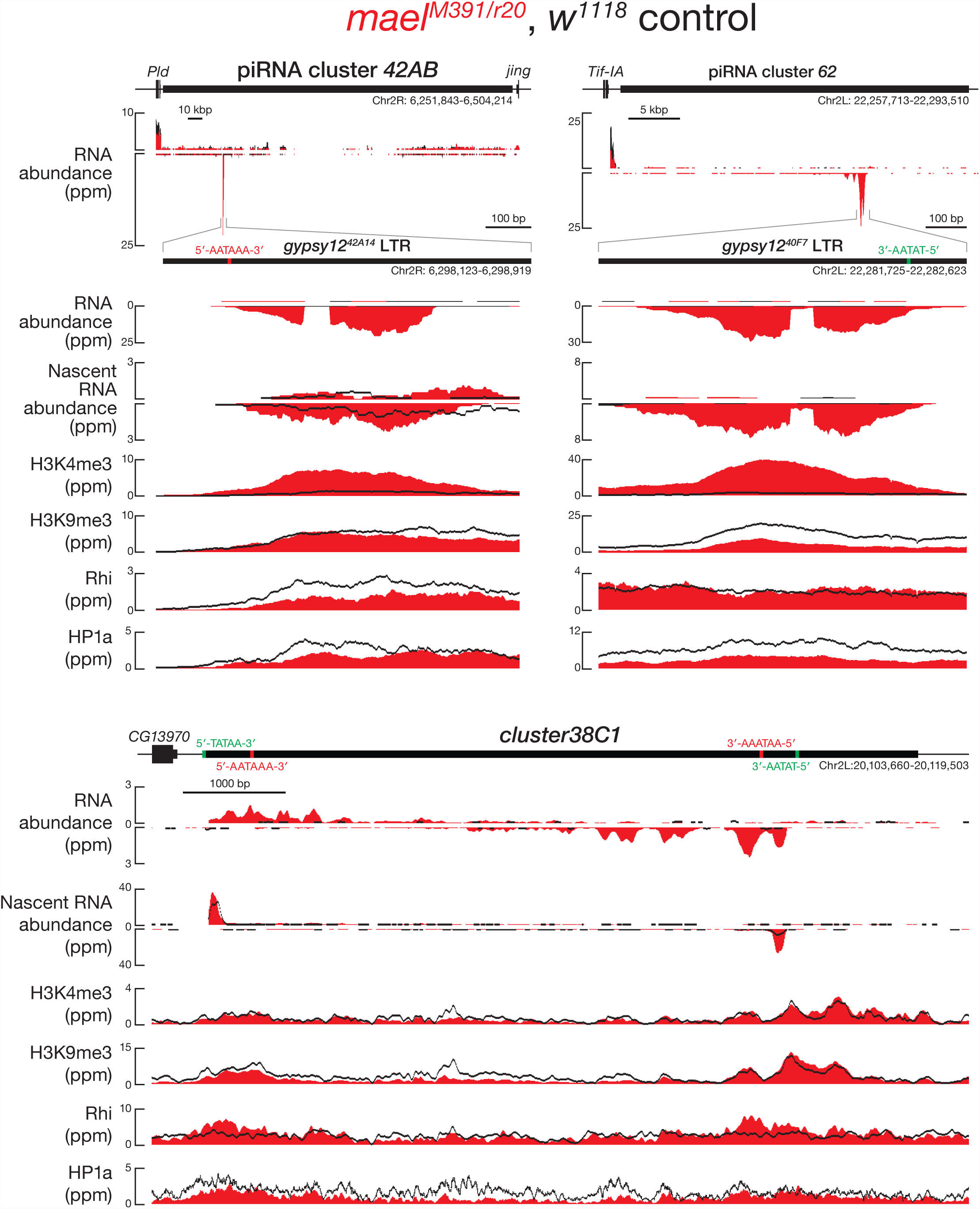
Canonical Transcription Initiation in Clusters *42AB, 62*, and *38C1* Without Mael. RNA abundance (RNA-seq), nascent RNA abundance (GRO-seq), and protein density for H3K4me3, H3K9me3, Rhi, and HP1a (ChIP-seq) at the dual-strand piRNA clusters *42AB*, and *cluster62*, and *38C1* was measured for wild-type control and *mael^M391/r20^*. Mutant ovaries are shown in red while wild-type controls are shown in black. Annotated transcription start (green) and termination sites (red) are labeled. Data are reads mapping uniquely to the genome from a representative experiment.

Loss of Mael also led to increased use of the canonical Pol II promoters flanking dual-strand *cluster38C1*. Unlike typical piRNA clusters, *cluster38C1* can sustain piRNA precursor production in mutants that disrupt incoherent transcription (Mohn et al., 2014; Chen et al., 2016; Andersen et al., 2017). In *mael^M391/r20^* ovaries, transcription initiating at canonical TATA-box sequences flanking the cluster increased for both the plus (mean increase *mael^M391/r20^*/control = 3 ± 1; *p* = 0.046) and minus (mean increase *mael^M391/r20^*/control = 5 ± 2; *p* = 0.014) genomic strands (Figure 1). Similarly, the steady-state abundance of *cluster38C1* RNA >150 nt long increased ∼15-fold in *mael^M391/r20^* ovaries (mean *mael^M391/r20^*/control = 15 ± 5; *p* = 3.1 × 10^−4^).

Unlike the *gypsy12* LTRs in *42AB* and *cluster62*, the density of the active chromatin mark H3K4me3 at *cluster38C1* was essentially unchanged in *mael^M391/r20^* ovaries (Figure 1). A combination of canonical and Rhi-dependent incoherent transcription has been proposed to produce piRNA precursor RNA from *cluster38C1* (Andersen et al., 2017). We speculate that because the wild-type level of H3K4me3 at *cluster38C1* already suffices to initiate transcription at the flanking promoters, no further increase occurs in *mael^M391/r20^* mutants.

### A Reporter for Transcription in Dual-Strand Clusters

piRNA clusters are highly repetitive, complicating bioinformatic analyses. The *P{GSV6}42A18* fly strain inserts a GAL4-responsive *gfp* transgene, containing five tandem repeats of UAS and a core promoter derived from the Hsp70Bb gene into *42AB* (Chendrimada et al., 2005), allowing *gfp* to be used as a proxy for canonical euchromatic transcription within piRNA clusters. Like *42AB* itself, *P{GSV6}42A18* requires Rhi, Piwi, and Armi to produce sense and antisense piRNAs (Figure S2A; Han et al., 2015a). In control flies, the *P{GSV6}42A18* transgene resembles the piRNA cluster in which it resides, with a high density of H3K9me3, HP1a, and Rhi across its sequence; expression of both *gfp* mRNA and protein was essentially undetectable even when the strong transcriptional activator GAL4-VP16 was co-expressed from the germline-specific *nanos* (*nos*) promoter (Figures 2A and 2B).

**Figure 2.**
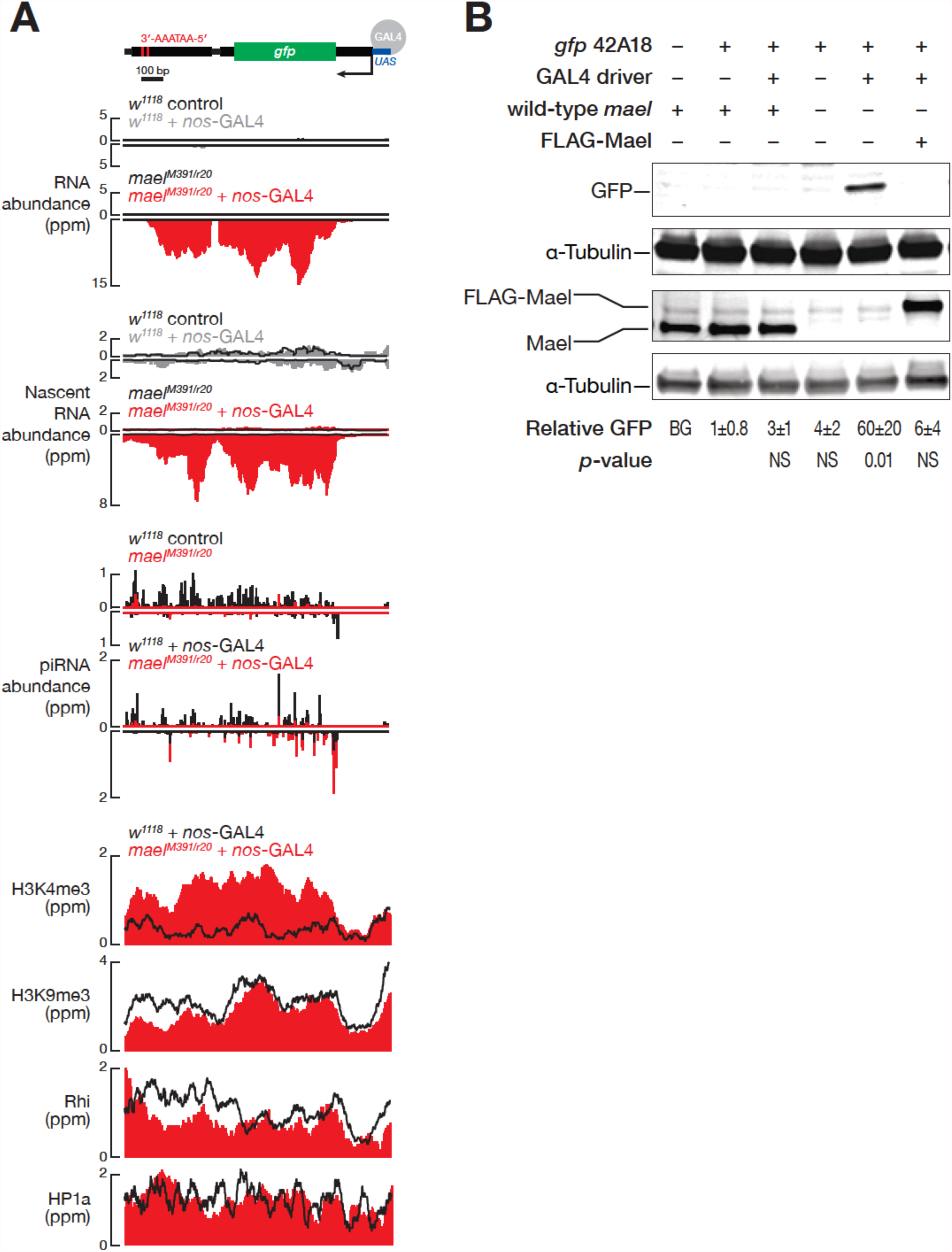
Mael Represses the Canonical Transcription of a Euchromatic Reporter in 42AB. (A) RNA-seq, GRO-seq, piRNA, and H3K4me3, H3K9me3, Rhi, and HP1a ChIP-seq profiles for *P{GSV6}42A18*, a *gfp* transgene inserted into *42AB* containing five tandem GAL4-binding upstream activating sequences with a core promoter derived from the *Hsp70Bb* gene (Chendrimada et al., 2005). For clarity, only one UAS is depicted here and in Figures 5B and S2A. The *gfp* transgene contains an intron and canonical poly(A) sites in the 3′ UTR. Reads from *mael^M391/r20^* mutants are shown in red while wild-type controls are shown in black. Annotated TTSs (red) are labeled. RNA-, GRO-, and piRNA data are for uniquely mapping reads from the mean of three biological samples. ChIP-seq data are for uniquely mapping reads from the mean of two biological samples. (B) Representative Western blots for GFP, Mael, or α-Tubulin (α-Tub) from ovaries with the genotype given below. GFP Western signal is the mean of three biological replicates normalized to α-Tub and is given in arbitrary units. *p*-values were measured using an unpaired, two-tailed t-test compared to *^w1118;^ P{GSV6}42A18*/+; +. Uncropped gel images can be found in Figure S2D.

In contrast, in *mael^M391/r20^* mutant ovaries, the *P{GSV6}42A18* transgene driven by GAL4-VP16 produced correctly spliced *gfp* mRNA that terminated at a canonical polyadenylation signal sequence; the appearance of *gfp* mRNA was accompanied by increased transcription (mean *mael^M391/r20^*/control = 80 ± 60; *p* = 3.0 × 10^−3^) and H3K4me3 (>3-fold increase) across the *gfp* transgene (Figure 2A). Moreover, the *gfp* mRNA in *mael^M391/r20^* mutants was translated into full-length GFP protein (Figures 2B and S2B). Finally, a transgene encoding FLAG-Mael restored repression of *gfp* in *mael^M391/r20^*, demonstrating that loss of Mael, and not a secondary mutation, caused inappropriate GFP expression from the transgene inserted into *42AB* (Figure 2B). Together, our data suggest that Mael represses canonical, promoter driven Pol II transcription in dual-strand piRNA clusters.

### Many Normally Repressed Pol II Promoters are Activated in *mael* Mutant Ovaries

Without Mael, RNA accumulates from both individual euchromatic transposons outside clusters (Sienski et al., 2012; Muerdter et al., 2013) and heterochromatic transposon sequences within clusters (Figures S1 and S3A). To further test the idea that Mael represses canonical transcription at sites of Rhi-driven incoherent transcription, we examined in more detail those transposons whose steady-state RNA abundance increased in *mael^M391/r20^* ovaries. Overall, steady-state RNA from 410 individual transposons within piRNA clusters and 1075 transposons outside clusters at least doubled in *mael^M391/r20^* ovaries (FDR ≤ 0.05; Figure S3A). Among these derepressed transposons, 182 overlapped H3K4me3 peaks whose area more than doubled in *mael* mutants (70 within and 112 outside clusters; Figure S3A). Moreover, of these 182 transposon loci, spliced transcripts—measured by the abundance of uniquely mapping exon-exon junction RNA-seq reads—more than doubled for 29 (13 within and 16 outside clusters) in *mael^M391/r20^* ovaries (Figure S3A).

In flies, most promoters are uni-directional. Therefore, antisense reads are unlikely to be products of canonical promoter-dependent transcription (Nechaev et al., 2010). In *mael^M391/r20^* mutants, antisense transposon transcripts within and outside piRNA clusters also increased for 427 loci (153 within and 274 outside clusters). Of these, 39 had overlapping H3K4me3 peaks (19 within and 20 outside clusters). Among these 39 antisense transposon loci, exon-exon junction reads established that the transcripts from at least 13 were spliced (6 within and 7 outside clusters; Figure S3A). Although sense transposon RNA abundance was ∼10-fold higher than antisense, the more than twofold increase in spliced antisense transcripts with overlapping H3K4me3 peaks suggests that cryptic antisense promoters may be active in *mael^M391/r20^* ovaries.

Consistent with an increase in canonical transcription, we also detected a fourfold increase in nascent transcripts from piRNA clusters (Figure S1). Transcription increased at 3,798 individual transposon loci both within (932) and outside (2,866) piRNA clusters in *mael^M391/r20^* ovaries (Figure S3B). Of these loci, 100 within and 184 outside piRNA clusters were also associated with increased H3K4me3 peaks (Figure S3B). Consistent with our RNA-seq results, we also found transposons that were actively transcribed in the antisense orientation that overlapped with H3K4me3 peaks in *mael^M391/r20^* mutants (Figure S3B). We conclude that loss of Mael increases canonical Pol II transcription from both euchromatic transposons outside piRNA clusters and from heterochromatic transposons within piRNA clusters.

### Heterochromatin is Largely Intact in *mael* Mutant Ovaries

Does loss of heterochromatin at transposons explain the increase in their transcription in *mael^M391/r20^* ovaries? We used chromatin immunoprecipitation sequencing (ChIP-seq) to examine the density of H3K9me3, HP1a, and Rhi in wild-type and *mael^M391/r20^* ovaries. Without Mael, we observed a modest (1.4–2.8-fold) decrease in the repressive heterochromatin mark H3K9me3 at the *gfp* insertion in *42AB*, *gypsy12^cluster62^*, *cluster38C1*, and *gypsy12^42AB^* (Figures 1 and 2A). HP1a binds H3K9me3, compacts chromatin, and, like Rhi, decorates piRNA clusters (Bannister et al., 2001; Jacobs and Khorasanizadeh, 2002; Nielsen et al., 2002; Vermaak and Malik, 2009; Klenov et al., 2014). Like H3K9me3, we only saw small changes in HP1a or Rhi at gypsy12*^42AB^*, gypsy12*^cluster62^*, cluster38C1, and gfp (Figures 1 and 2A).

Despite retaining the hallmarks of heterochromatin, gypsy12*^42AB^*, gypsy12*^cluster62^*, and the gfp transgene, all acquired the active transcription mark H3K4me3 when they became transcribed in mael*^M391/r20^* mutant ovaries (Figures 1 and 2A). We note that the coexistence of active and repressive chromatin marks is found at active genes embedded in heterochromatin (Riddle et al., 2011). We conclude that failure to repress canonical transcription of piRNA clusters in mael mutants does not reflect a loss of heterochromatin.

Moreover, loss of Mael had no detectable effect on heterochromatin elsewhere in the genome. We divided the genome into 1 kb bins and examined the densities of H3K9me3, HP1a, and Rhi across piRNA clusters, transposons, protein-coding genes, and other genomic regions in mael*^M391/r20^* and control ovaries. Across the genome, the median density of H3K9me3, HP1a, or Rhi in any category did not change by more than twofold between *mael ^M391/r20^* and control (Figure 3). We conclude that Mael is not required to maintain H3K9me3 in the germline. Similar results have been reported for Mael in the follicle cell-like, cultured ovarian somatic cell line (Sienski et al., 2012).

**Figure 3.**
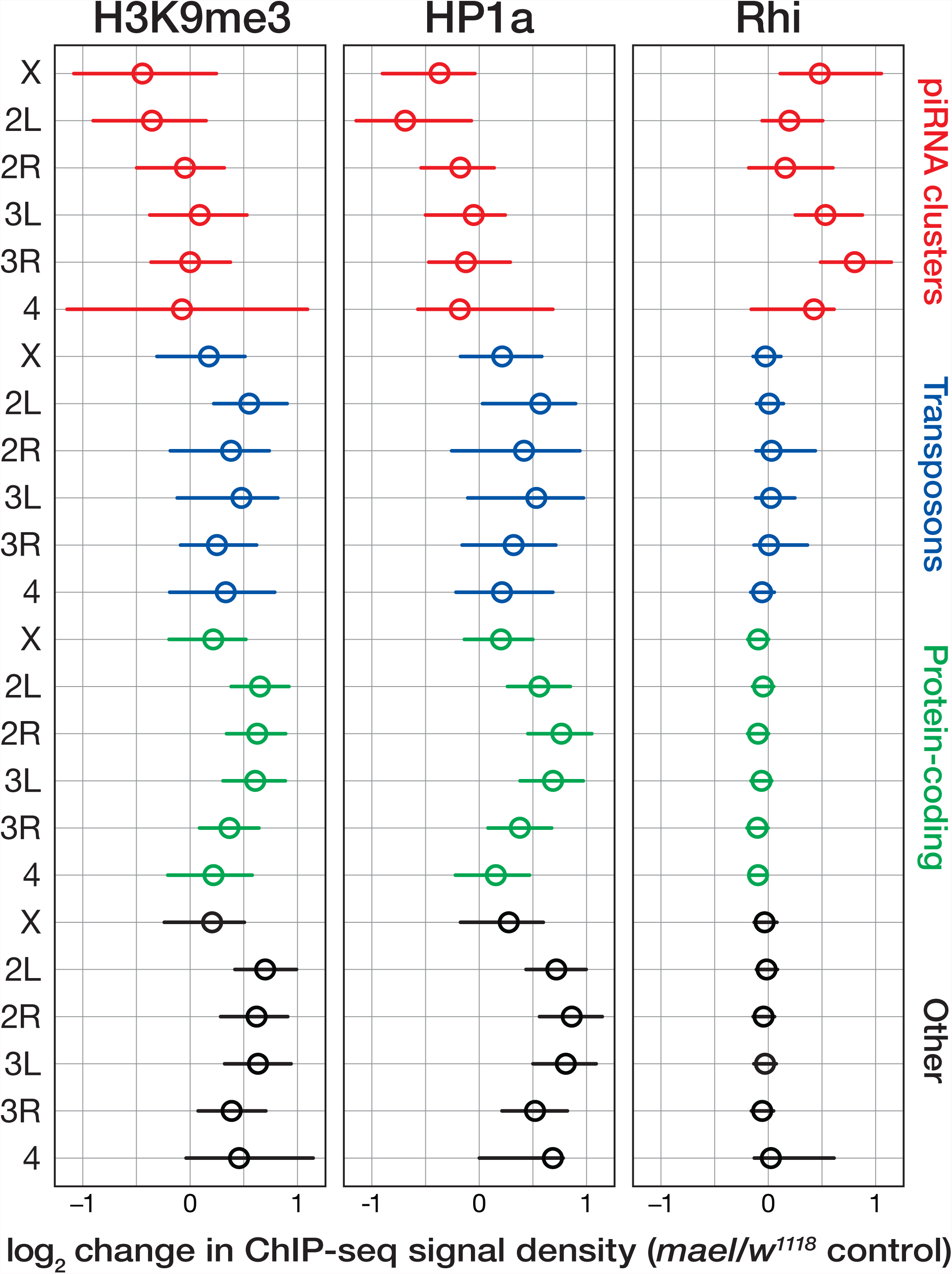
Loss of Mael does not alter Heterochromatin. ChIP-seq was used to measure the change (*mael*/control) between *mael^M391/r20^* and control ovaries in the density of H3K9me3, HP1a, or Rhi for piRNA clusters, transposons, and protein-coding genes. Circle denotes the median. Left and right whiskers encompass the first to third quartiles. Density was calculated using a 1 kbp sliding window with a 500 bp step and was normalized to the total number of reads mapping to each chromosome arm. Data are for uniquely mapping reads from two biologically independent samples.

### *mael* Mutants Produce Fewer piRNAs

Although nascent and steady-state RNA abundance from piRNA clusters increased in *mael^M391/r20^* mutant ovaries, piRNA abundance decreased (all piRNAs, *mael^M391/r20^*/control = 0.39 ± 0.03, *p* = 6.8 × 10^−6^; uniquely mapping piRNAs, *mael^M391/r20^*/control = 0.28 ± 0.02, *p* = 9.4 × 10^−7^; *n* = 3; Figure 4A). The abundance of piRNAs from *gypsy12^42AB^* (*mael^M391/r20^*/control = 0.060 ± 0.002, *n* = 3; *p* = 7.0 × 10^−9^), *gypsy12^cluster62^* (*mael^M391/r20^*/control = 0.030 ± 0.003, *n* = 3; *p* = 4.1 × 10^−6^), the *gfp* transgene inserted into 42AB (*mael^M391/r20^/*control = 0.09 ± 0.01, *n* = 3; *p* = 3.56 × 10^−7^), as well as from cluster38C1 (*mael^M391/r20^*/control = 0.10 ± 0.01, *n* = 3; *p* = 5.92 × 10^−7^), were all more than 10 times lower in *mael^M391/r20^* ovaries (Figures 2A and 4B). The loss of piRNAs was particularly acute for dual-strand piRNA clusters. For example, piRNA abundance decreased 3–5-fold for the paradigmatic uni-strand clusters *cluster2* (*mael^M391/r20^*/control = 0.21 ± 0.02; *p* = 8.4 × 10^−6^) and *flam* (*mael^M391/r20^*/control = 0.33 ± 0.03; *p* = 1.8 × 10^−5^), but 32-fold for the dual-strand cluster *42AB* (*mael^M391/r20^*/control = 0.053 ± 0.005; *p* = 9.7 × 10^−10^) and 7.6-fold for *80EF* (*mael^M391/r20^*/control = 0.13 ± 0.01; *p* = 4.6 × 10^−7^). piRNA abundance also declined for telomeric clusters (*mael^M391/r20^/*control = 0.031 ± 0.001; *p* = 1.0 × 10^−8^). These data suggest that the canonically transcribed RNA produced from dual-strand clusters in the absence of Mael cannot efficiently enter the piRNA biogenesis pathway.

**Figure 4.**
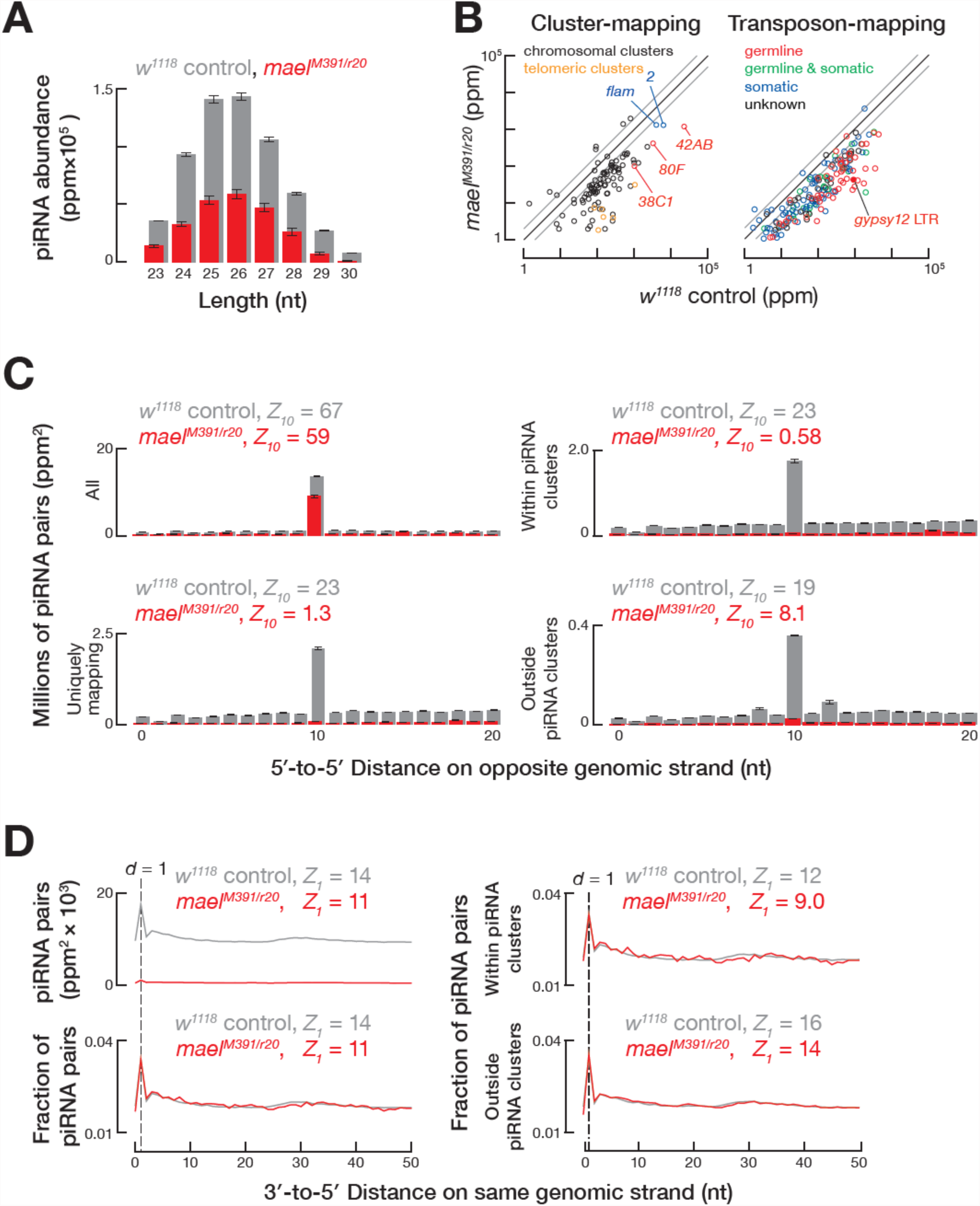
Fewer piRNAs in *mael^M391/r20^* Mutants. (A) Length distribution of piRNAs from *^w1118^* control and *mael^M391/r20^* ovaries. (B) Scatter plot comparing piRNAs that uniquely map to germline (red), soma (blue), intermediate (green), or unknown (black) transposons. Labeled uni-strand clusters are depicted in blue while labeled dual-strand clusters are depicted in red. Telomeric piRNA clusters are displayed as orange circles. piRNAs mapping to *gypsy12* LTRs are labeled and shown as a solid red circle. The grey line signifies a ≥2-fold change. Data are from three biologically independent samples. (C) Ping-pong analysis for all or uniquely mapping total piRNAs. Data are mean ± S.D. (*n* = 3). (D) Phasing analysis for the distances from the 3′ ends of upstream piRNAs to the 5′ ends of downstream piRNAs on the same genomic strand. Data are for uniquely mapping reads. All data are mean ± S.D. (*n* = 3).

The decreased abundance of piRNAs from dual-strand clusters in *mael^M391/r20^* mutants was accompanied by a marked loss of piRNA amplification (Figure 4C). This defect, however, did not reflect a loss of the ping-pong machinery (*mael^M391/r20^*, *Z_10_* = 59; control, *Z_10_* = 67). However, *mael^M391/r20^* ovaries showed no significant ping-pong among piRNAs unambiguously mapping to dual-strand clusters (*mael^M391/r20^*, *Z_10_* = 0.58; control, *Z_10_* = 23). In contrast, piRNAs mapping outside clusters continued to be amplified (*mael^M391/r20^*, *Z_10_* = 8.1; control, *Z_10_* = 19; Figure 4C).

Like secondary piRNA production by the ping-pong cycle, the machinery required to produce phased primary piRNAs appears unaltered in *mael^M391/r20^* mutants. Despite the reduced abundance of piRNA in *mael^M391/r20^* ovaries, significant tail-to-head primary piRNA phasing remained (all piRNAs: *mael^M391/r20^*, *Z_1_* = 11 versus control, *Z_1_* = 14; Figure 4D). Unlike ping-pong amplification, cluster-mapping piRNAs continued to be produced by the phased primary piRNA pathway (within clusters: *mael^M391/r20^*, *Z_1_* = 9.0 versus control, *Z_1_* = 12; outside clusters: *mael^M391/r20^*, *Z_1_* = 14 versus control, *Z_1_* = 16; Figure 4D). We conclude that the primary piRNA biogenesis machinery is intact in the absence of Mael.

### Armi and Piwi, but not Rhi, Repress Transcription in Dual-Strand Clusters

The current model for piRNA biogenesis in flies places Armi upstream and Mael downstream of Piwi (Malone et al., 2009; Haase et al., 2010; Saito et al., 2010; Sienski et al., 2012; Czech et al., 2013; Pandey et al., 2017; Rogers et al., 2017). Consistent with this model, loss of either Armi or nuclear Piwi phenocopies the loss of Mael. For example, cluster-mapping steady-state transcript abundance increased to similar levels in *armi^72 1/G728E^*, *piwi^2/Nt^*, and *mael^M391/r20^* ovaries compared to control (Figure S4A). The abundance of RNA from *cluster38C1*, *gypsy12^42AB^*, and *gypsy12^cluster62^*, increased in all three mutants. Moreover, the RNA produced from both the *gypsy12^42AB^* and *gypsy12^cluster62^* LTRs, was spliced, consistent with a failure to repress canonical transcription. Loss of Piwi or Armi similarly increased the steady-state abundance of RNA or protein from the UAS/Gal4-driven *gfp* transgene inserted into piRNA cluster *42AB* (Figures 5B and 5C). Finally, *piwi^2/Nt^*, *armi^72 1/G728E^*, and *mael^M391/r20^* mutant ovaries all had fewer *gfp*-mapping piRNAs than control (Figure S2A; Han et al., 2015a). Our results suggest that Armi, Piwi, and Mael act in a common pathway to repress canonical transcription in dual-strand piRNA clusters.

**Figure 5.**
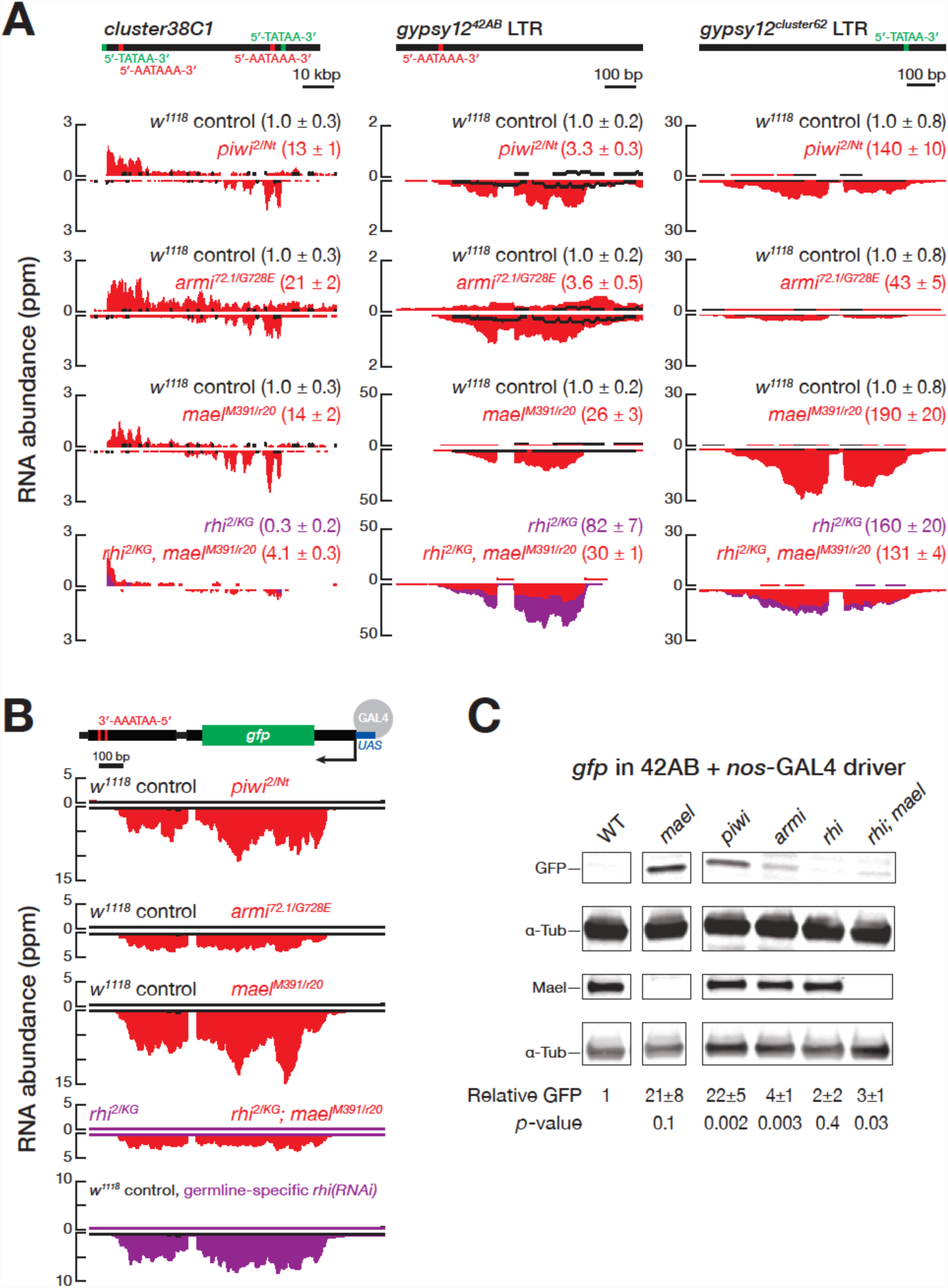
Armi and Piwi, but not Rhi, are Required to Repress Canonical Transcription in Dual-Strand Clusters. (A) RNA-seq profiles for *gypsy12^42AB^* (left), *gypsy12^cluster62^* (center), *cluster38C1* (right), and (B) *P{GSV6}42A18* from control, *armi^72 1/G728E^*, *piwi^2/Nt^*, *rhi^2/KG^*, *mael^M391/r20^*, and *rhi^2KG^; mael^M391/r20^*, and germline-specific *rhi(RNAi)* ovaries. (C) Representative Western blots and quantifications for GFP, Mael, or α-Tubulin (α-Tub) from ovaries. Uncropped gel images can be found in Figure S2D.

Our data also suggest that by promoting incoherent transcription in otherwise repressive heterochromatin, Rhi creates the requirement for Mael to suppress canonical transcription of *gypsy12* in dual-strand piRNA clusters. In *rhi^2/KG^* ovaries, steady-state, spliced RNA initiating in the *gypsy12^42AB^* and *gypsy12^cluster62^* LTRs increased >80- and >160-fold, respectively, despite the presence of normal levels of Mael (Figure 5A). Removing both Rhi and Mael (*rhi^2/KG^*, *mael^M391/r20^*) did not appreciably increase the abundance of spliced transcripts from the two *gypsy12* LTRs (Figure 5A), consistent with the idea that without Rhi, Mael fails to repress canonical transcription from the *gypsy12* LTRs. In contrast, loss of Rhi did not produce detectable mRNA or protein from the *gfp* transgene inserted in *42AB* (Figures 5B and 5C).

Loss of Rhi alone leads to canonical transcription at the two *gypsy12* LTRs, yet canonical transcription of the *gfp* transgene in 42AB requires the loss of Mael and the presence of Rhi. Why does *gfp* transgene transcription require Rhi? Perhaps Rhi creates a chromatin environment that is transcriptionally permissive for euchromatic genes but repressive for heterochromatic genes. We speculated that low levels of Rhi might suffice to support canonical transcription of the *gfp* transgene, even in the presence of Mael. To test this idea, we used RNAi to reduce, but not eliminate, Rhi in the ovary germline (Figures S4B and S4C). Consistent with the hypothesis, *rhi(RNAi)* ovaries produced spliced *gfp* mRNA, unlike *rhi^2/KG^* null mutants (Figure 5B). We propose that by promoting incoherent transcription, Rhi also promotes canonical transcription, which Mael then acts to repress.

### Maelstrom is a Suppressor of Position-Effect Variegation

Position-effect variegation (PEV) is the phenomenon in which gene expression is dependent on chromatin context. Suppressors of PEV are often genes involved in making or maintaining heterochromatin, such as *Suppressor of variegation 205* (*Su(var)2-5*), which encodes HP1a (Eissenberg et al., 1992). The loss of *gfp* silencing in *mael^M391/r20^* mutants is genetically similar to a loss-of-function mutation in a suppressor of PEV. In fact, the transgene *P{GSV6}42A18*, which introduced *gfp* into *42AB*, also contains the marker gene *w^+mc^* (mini-*white*). Like the *gfp* transgene, *w^+mc^* was silenced: *w^1118^*; *P{GSV6}42A18* flies have white eyes. Intriguingly, the eyes of *w^1118^*; *P{GSV6}42A18; mael^M391/r20^* flies were orange (Figure 6).

**Figure 6.**
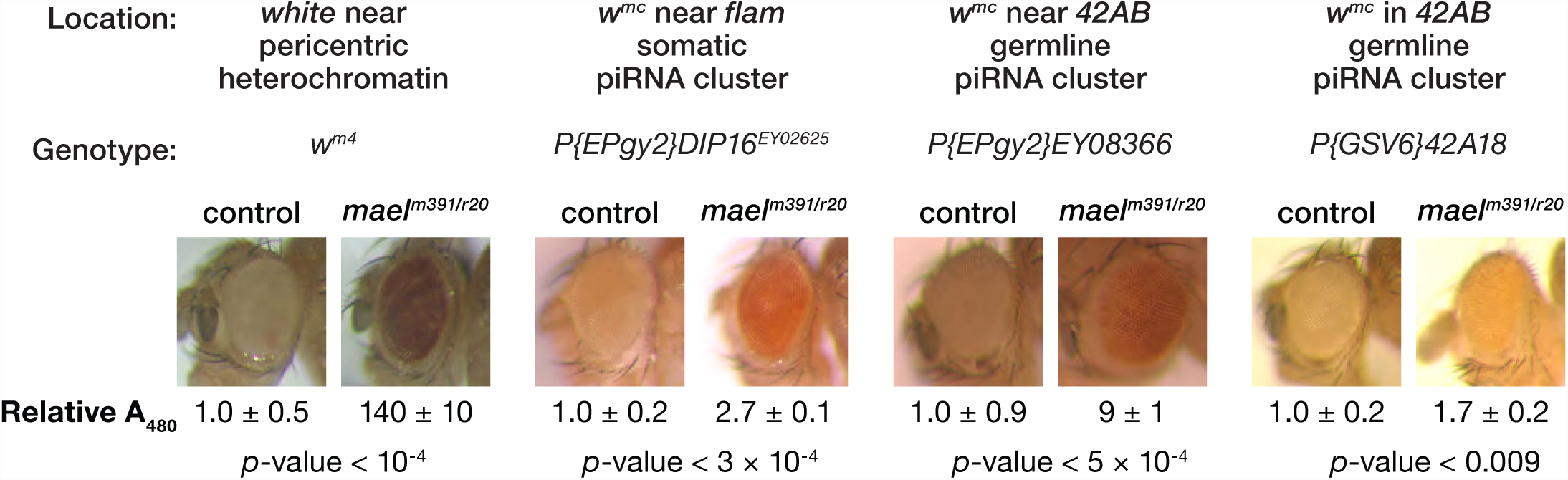
Mael Represses Heterochromatic Sequences in the Soma. Representative images of eyes from *mael^M391/r20^* and control flies from different variegating strains. The source of *white* is shown above the image. Red pigment was quantified by measuring the absorbance at 480 nm (A480) of pigment extracted with acidic ethanol. Data are mean ± S.D. (*n* = 3).

To further explore a potential role for *mael* in PEV, we employed two additional transgenic PEV models: *P{EPgy2}EY08366*, which inserts *w^+mc^* near *42AB*, and *P{EPgy2}DIP16^EY02625^*, which inserts *w^+mc^* near *flam*. PEV at both reporters has been shown to be suppressed by *Su(var)2-5* or *Su(var)3-9* mutations, but not by *piwi* or *aub* mutations (Moshkovich and Lei, 2010). In contrast, loss of Mael significantly increased red pigment expression for both reporters (Figure 6).

Finally, we tested the effect of loss of Mael on the classic PEV allele, *white-mottled4* (*w^m4^*), in which a chromosomal inversion moves the euchromatic *white* gene near the centromeric heterochromatin (Muller, 1930). *w^m4^* flies have white eyes. As expected, both *Su(var)2-5* and *Su(var)3-9* heterozygotes restored red eye color to *w^m4^* flies, increasing the amount of red pigment 120- and 150-fold, respectively (Table S1). Similarly, loss of one copy of *mael* also suppressed *white* silencing, increasing red pigmentation 80–90 times (Figure 6 and Table S1). Thus, *mael* is a classical suppressor of PEV. We suggest that *mael* mutants suppresses PEV in the somatic cells of the adult eye by the same mechanism it suppresses *gfp* and *gypsy12* expression in dual-strand, germline piRNA clusters, by repressing canonical euchromatic transcription in heterochromatic genomic regions.

## DISCUSSION

### Mael Represses Canonical Transcription Activated by Rhi

Fly piRNA clusters must solve a gene-expression paradox. They record the ancient and contemporary exposure of the animal to transposon invasion, and this information must be copied into RNA in order to generate protective, anti-transposon piRNAs. However, more recent transposon insertions retain the ability to produce mRNA encoding proteins required for their transposition. Recent evidence suggests that, in flies, dual-strand piRNA clusters solve this expression paradox by using Rhi to initiate incoherent transcription of unspliced RNA, the precursor to piRNAs, while repressing promoter-initiated, canonical transcription (Le Thomas et al., 2014; Mohn et al., 2014; Zhang et al., 2014; Andersen et al., 2017). Our data suggest that Mael is required for this second process, thereby allowing dual-strand piRNA clusters to safely generate piRNA precursor transcripts without risking production of transposon mRNAs (Figure 7). Notably, both Rhi and Mael require the nuclear protein Piwi and the piRNA biogenesis protein Armi for their function, suggesting that both the initiation of incoherent transcription by Rhi and the suppression of coherent transcription by Mael are directly or indirectly guided by piRNAs.

**Figure 7.**
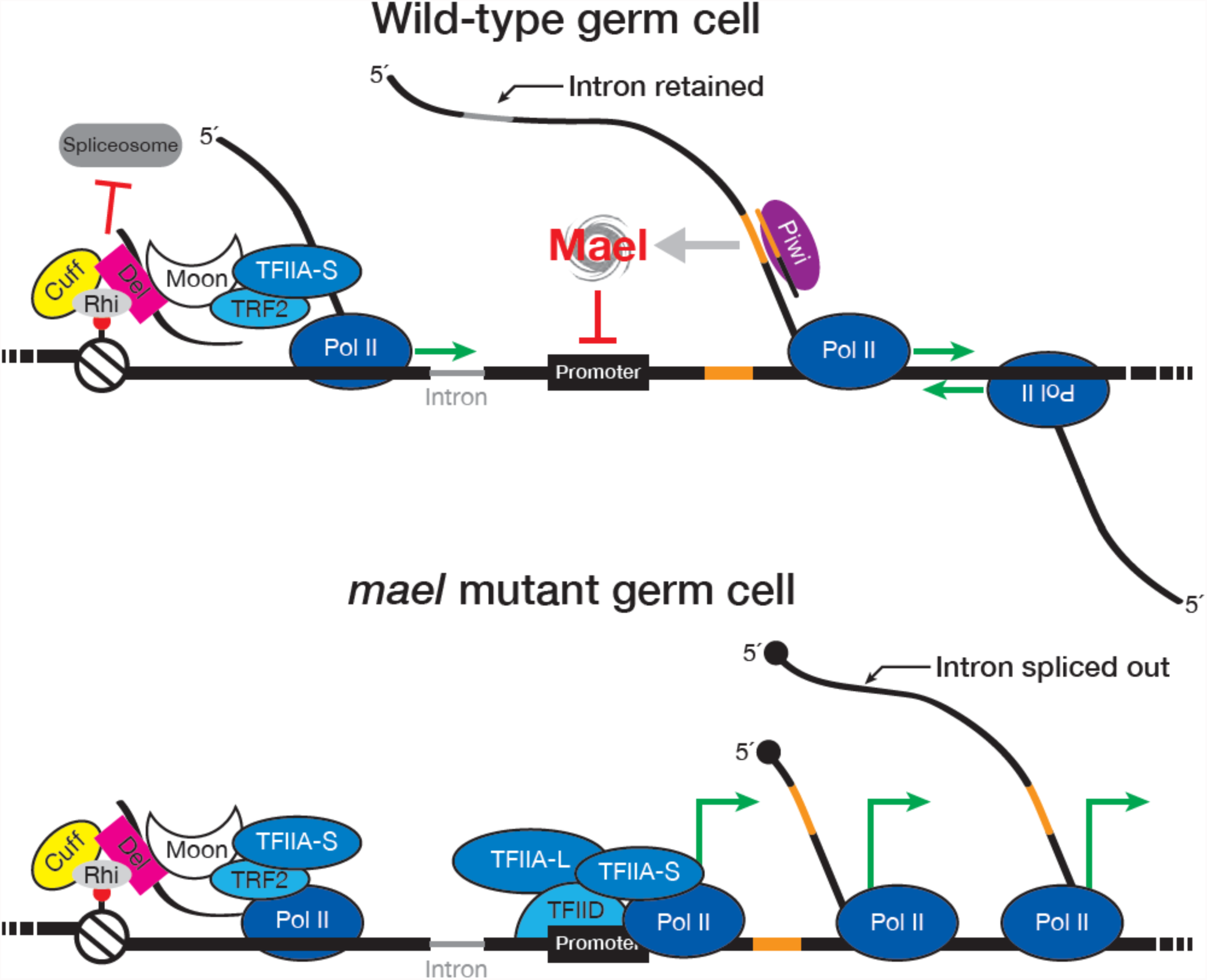
Model for Mael-Dependent Repression of Coherent Transcription. Dual-strand piRNA cluster transcription is dependent on Rhi and associated proteins, which allow Pol II to initiate independent of promoters and produce non-canonical transcripts. While non-canonical and “incoherent,” Rhi-mediated transcription may expose promoters that would otherwise be hidden by heterochromatin. Piwi, guided by piRNAs, binds to nascent transcripts and targets different proteins to genomic loci to repress transcription. One of these proteins is Mael, which blocks promoter-driven transcription. Without Mael, canonical transcripts are produced from dual-strand clusters which can be translated into protein.

### Mael is Required for Dual-Strand piRNAs

Although Mael acts after piRNA-guided Piwi, *mael^M391/r20^* ovaries have fewer piRNAs than control (Figures 4A and 4B). In particular, piRNAs mapping to dual-strand clusters, including *P{GSV6}42A18*, severely decreased, despite a concomitant increase in canonical transcription from these same DNA loci. Our data suggest that in *mael^M391/r20^* mutants, Rhi-mediated incoherent transcription, cluster transcript export, and ping-pong amplification become uncoupled. Thus, instead of fueling piRNA production, the coherent transcripts from dual-strand piRNA clusters produced in the absence of Mael are translated into protein (Figures 2A and 2B).

### A Putative Conserved Role for Mael

Although Rhi is only expressed in germ cells, we find that *mael* is a suppressor of PEV. Although Piwi is not readily detectable outside the gonads, depleting Piwi in early embryos suppresses the variegation of a somatic reporter in adult flies (Gu and Elgin, 2013). Perhaps, Piwi guides Mael to somatic targets early in development (Akkouche et al., 2017), establishing heterochromatin that is maintained through adulthood. *D. melanogaster* does not have a somatic piRNA pathway outside the gonads, but most arthropods—including spiders (*Parasteatoda tepidariorum*), centipedes (*Strigamia maritima*), and insects (e.g., *Drosophila virilis*, *Aedes aegypti*, and *Trichoplusia ni*) have both somatic and germline piRNAs (Fu et al., 2018b; Lewis et al., 2018). Supporting this view, 12 arthropods with somatic piRNAs—including insects, spiders, horseshoe crabs, and scorpions—all express *mael* mRNA in their soma, while three insects with no detectable piRNAs outside the gonads had low or undetectable somatic *mael* mRNA (Figure S5 and Table S2).

In male mice, loss of MAEL also leads to loss of piRNAs, germline transposon expression, and male sterility (Costa et al., 2006; Soper et al., 2008; Aravin et al., 2009; Castañeda et al., 2014). As in flies, loss of MAEL in mice does not trigger loss of heterochromatin: DNA methylation of L1 elements is unchanged (Aravin et al., 2009). Despite these similarities, fly Mael does not appear to interact with other piRNA pathway factors (Sato et al., 2011), while mouse MAEL colocalizes and associates with piRNA pathway proteins and piRNA cluster transcripts and transposons (Aravin et al., 2009; Castañeda et al., 2014). Current evidence from mouse suggests that MAEL is involved in the processing of piRNA precursors (Soper et al., 2008; Aravin et al., 2009; Castañeda et al., 2014). Thus, fly Mael and mouse MAEL may be orthologs with divergent mechanisms or, perhaps, a role for MAEL in repressing canonical transcription in mammals remains to be discovered.

## STAR METHODS

Detailed methods are provided in the online version of this paper and include the following:

- KEY RESOURCES TABLE
- CONTACT FOR REAGENT AND RESOURCE SHARING
- METHOD DETAILS

- *Drosophila* Stocks
- General Methods
- Western Blotting
- Immunohistochemistry and Microscopy
- Female Fertility Assay
- Eye Pigment Assay
- Construction and Analysis of sRNA-Seq Libraries
- Ping-Pong Analysis
- Phasing Analysis
- Construction and Analysis of RNA-Seq Libraries
- Construction and Analysis of ChIP-Seq Libraries
- Construction and Analysis of GRO-Seq Libraries

## SUPPLEMENTAL INFORMATION

Supplemental Information includes 5 figures and 2 tables and can be found with this article online at **http://XYZXYZXYZ**.

## AUTHOR CONTRIBUTIONS

T.H.C., E.M., Z.W., and P.D.Z. conceived and designed the experiments. T.H.C. performed the experiments. E.M., T.H.C., and I.G. analyzed the sequencing data. T.H.C., E.M., Z.W., and P.D.Z. wrote the manuscript.

## ACKNOWLEDGEMENTS

We thank William Theurkauf for the Ago3 antibody; Julius Brennecke for the Mael antibody; Toshie Kai for the FLAG-Mael rescue fly; Cindy Tipping and Alicia Boucher for fly husbandry, and members of the Weng and Zamore laboratories for help, advice, discussions, and comments on the manuscript. This work was supported in part by National Institutes of Health grants P01HD078253 to Z.W. and P.D.Z. and GM65236 to P.D.Z.

## SUPPLEMENTAL FIGURE TITLES AND LEGENDS

**Figure S1.**
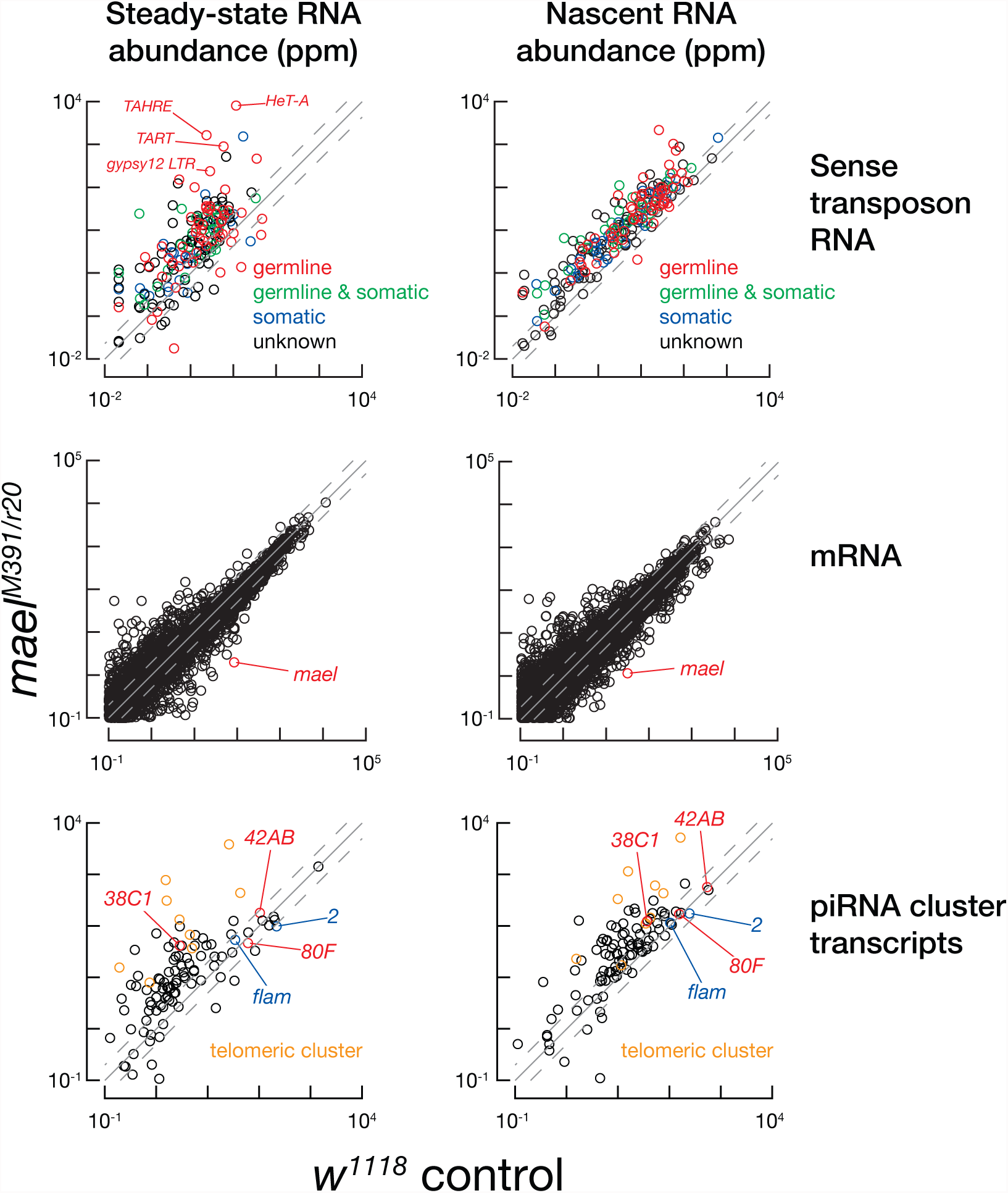
Scatter plots comparing the abundance of steady-state RNA (left) and nascent transcripts (GRO-seq; right) from *mael^M391/r20^* and control ovaries. Top, RNA to sense transposon sequences; middle, mRNA; bottom, RNA mapping to piRNA clusters. Transposons were classified according to their expression exclusively in the germline (red), both the germline and soma (green), or exclusively in the soma (blue) according to (Wang et al., 2015). Red circle: transcripts from the *mael* gene. Blue: uni-strand piRNA clusters. Red: dual-strand clusters. Orange: telomeric clusters are shown as circles. The data are for uniquely mapping reads from the mean of three independent biological samples.

**Figure S2, related to Figure 2.**
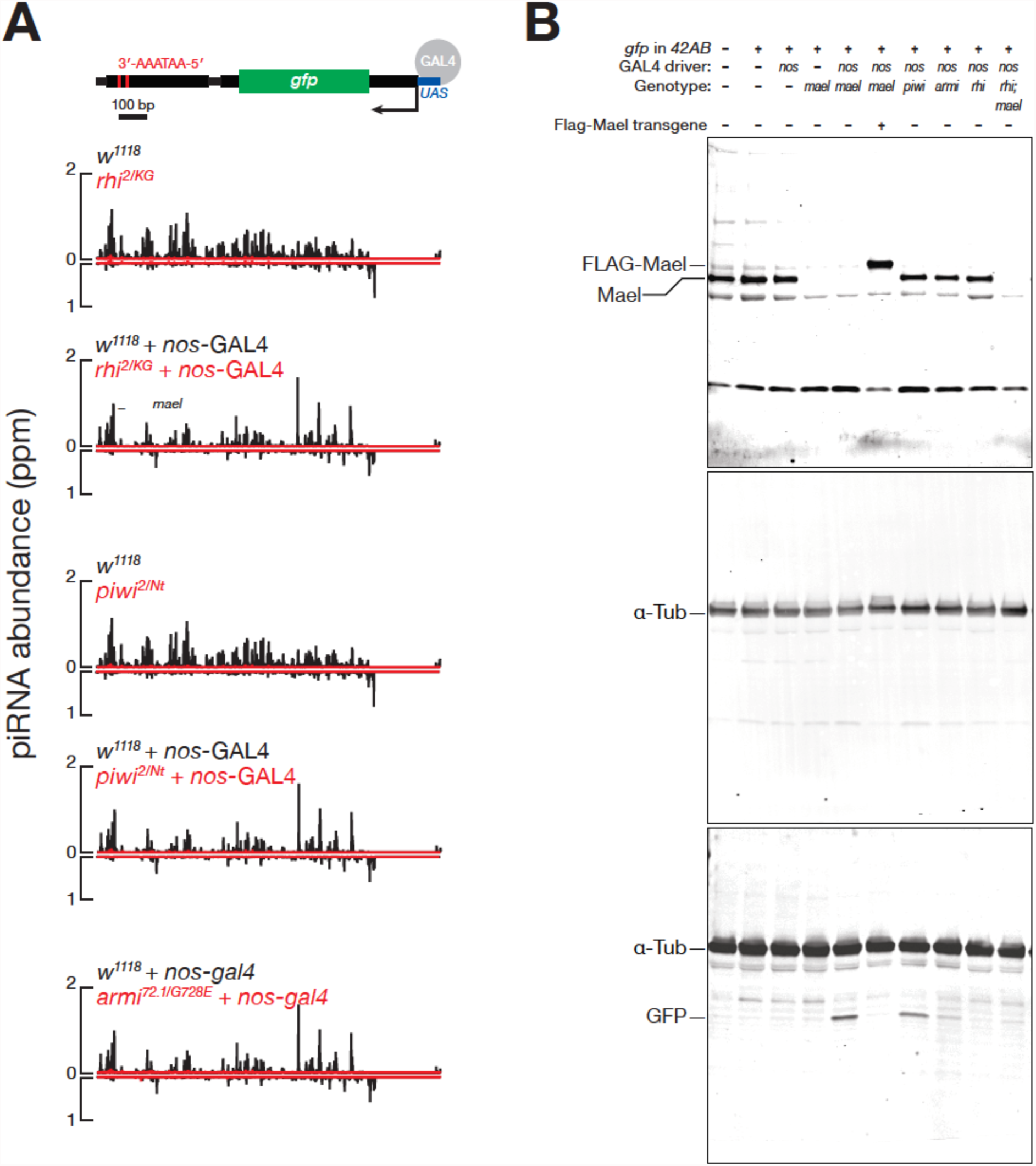
(A) Abundance of piRNAs mapping to *gfp* inserted into the 42AB piRNA cluster (*P{GSV6}42A18*) in *w^1118^* control, *rhi^2/KG^*, *piwi^2/Nt^*, and *armi^72 1/G728E^* ovaries. (B) Uncropped representative Western blots detecting GFP, Mael, and Tub.

**Figure S3, related to Figures 1 and 2.**
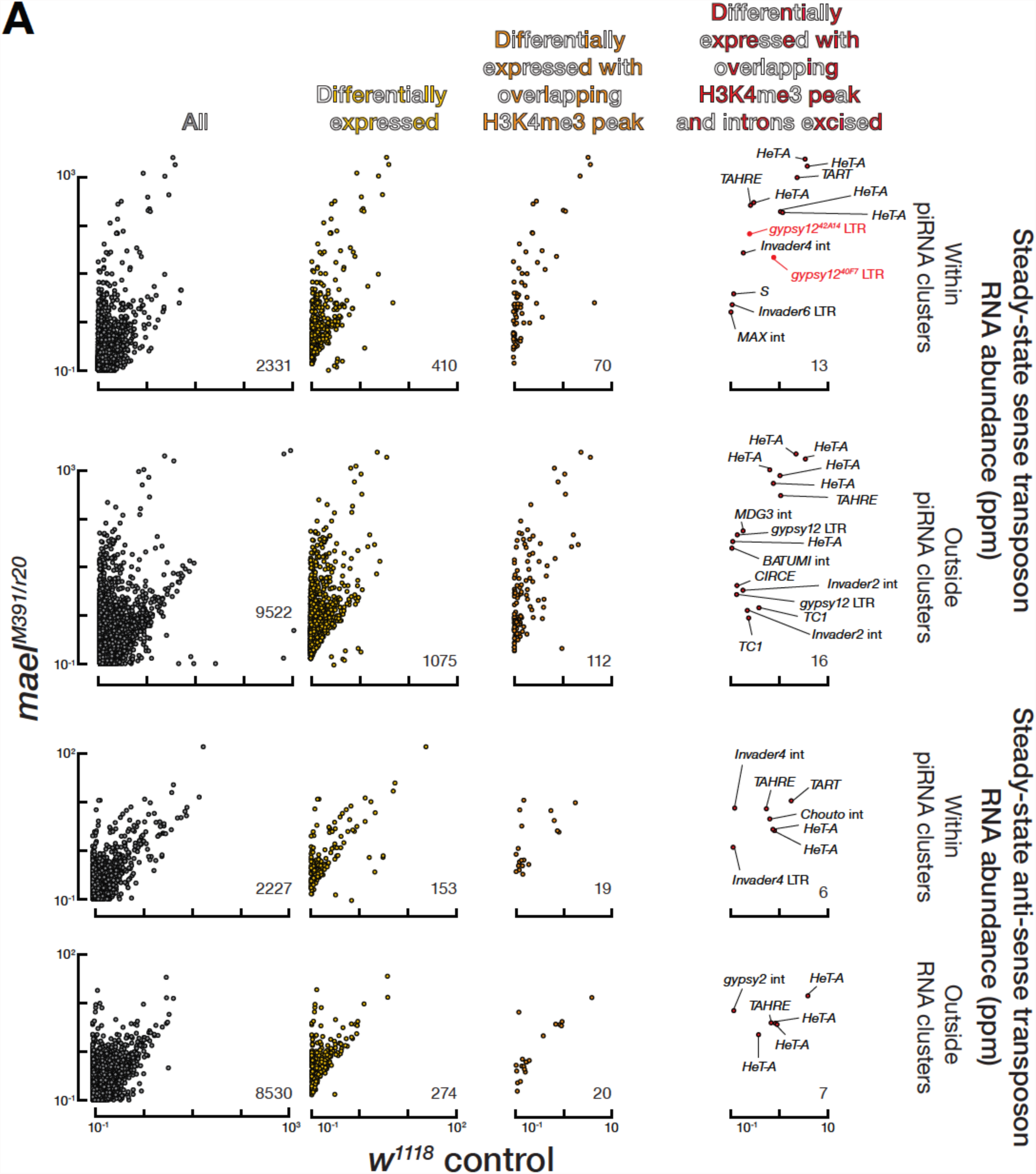
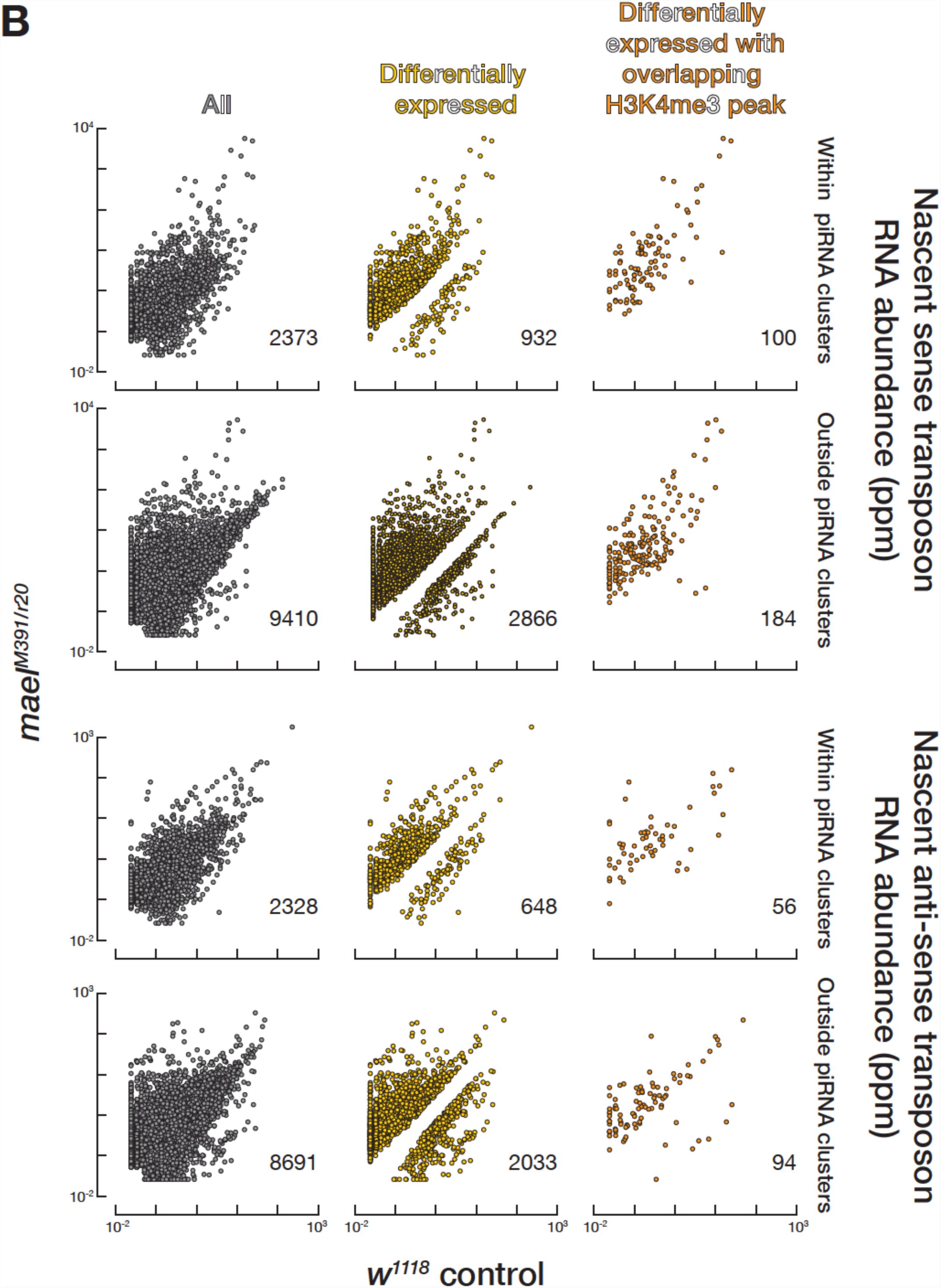
Scatter plots comparing (A) sense or antisense steady-state RNA (RNA-seq) or (B) sense or antisense nascent RNA (GRO-seq) abundance between *mael^M391/r20^* and control ovaries. Yellow: transposon loci differentially expressed between the two genotypes (≥2-fold increase, FDR ≤ 0.05). Orange: transposon loci both with overlapping H3K4me3 peaks and an increase in RNA in *mael^M391/r20^* mutants. Red, (A) only: transposon loci with differentially expressed, spliced transcripts (≥2-fold increase; FDR ≤ 0.05) and overlapping H3K4me3 peaks. The number of individual transposon loci is given in each scatter plot. All data are for uniquely mapping reads from the mean of six independent biological samples.

**Figure S4, related to Figure 4.**
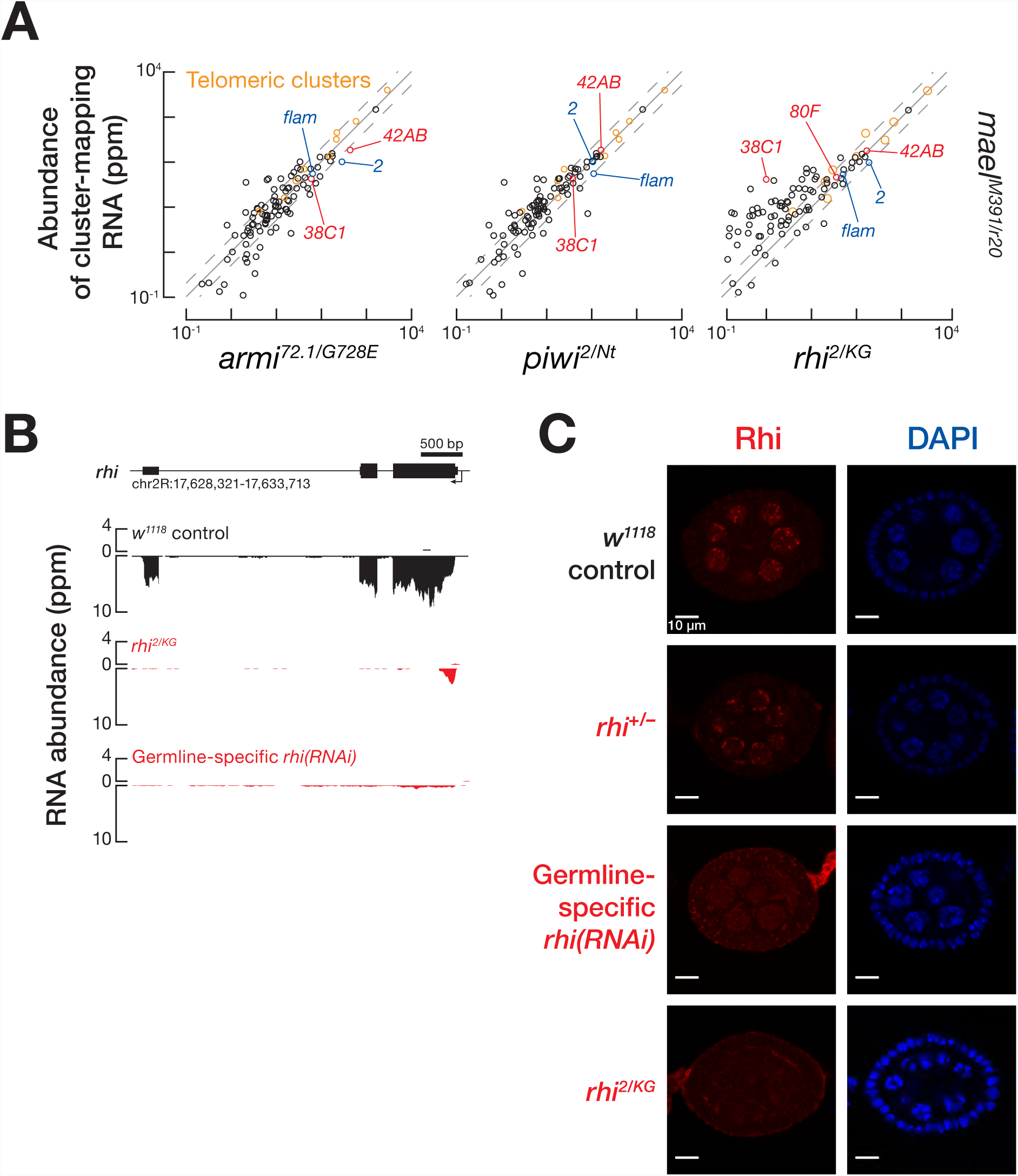
(A) Comparison of transcript (top) or piRNA abundance (bottom) for RNA mapping to piRNA clusters between *mael^M391/r20^* or *w^1118^* control and *armi^72 1/G728E^*, *piwi^2/Nt^*, or *rhi^2/KG^* mutant ovaries. Blue, uni-strand clusters; red, dual-strand clusters; orange, telomeric clusters. Hashed grey line, ≥2-fold change. Data are for uniquely mapping reads from the mean of three biological samples. (B) RNA abundance of *rhi* for *w^1118^* control, *rhi^2/KG^* mutant, and germline-specific *rhi(RNAi)* ovaries. (C) Immuno-detection of Rhi protein stage 4 egg chambers. DAPI, 4′,6-diamidino-2-phenylindole. (D) RNA abundance for *gfp* inserted into the 42AB piRNA cluster (*P{GSV6}42A18*) from germline-specific *rhi(RNAi)* and *w1118* control ovaries expressing germline GAL4-VP16.

**Figure S5, related to Figure 1.**
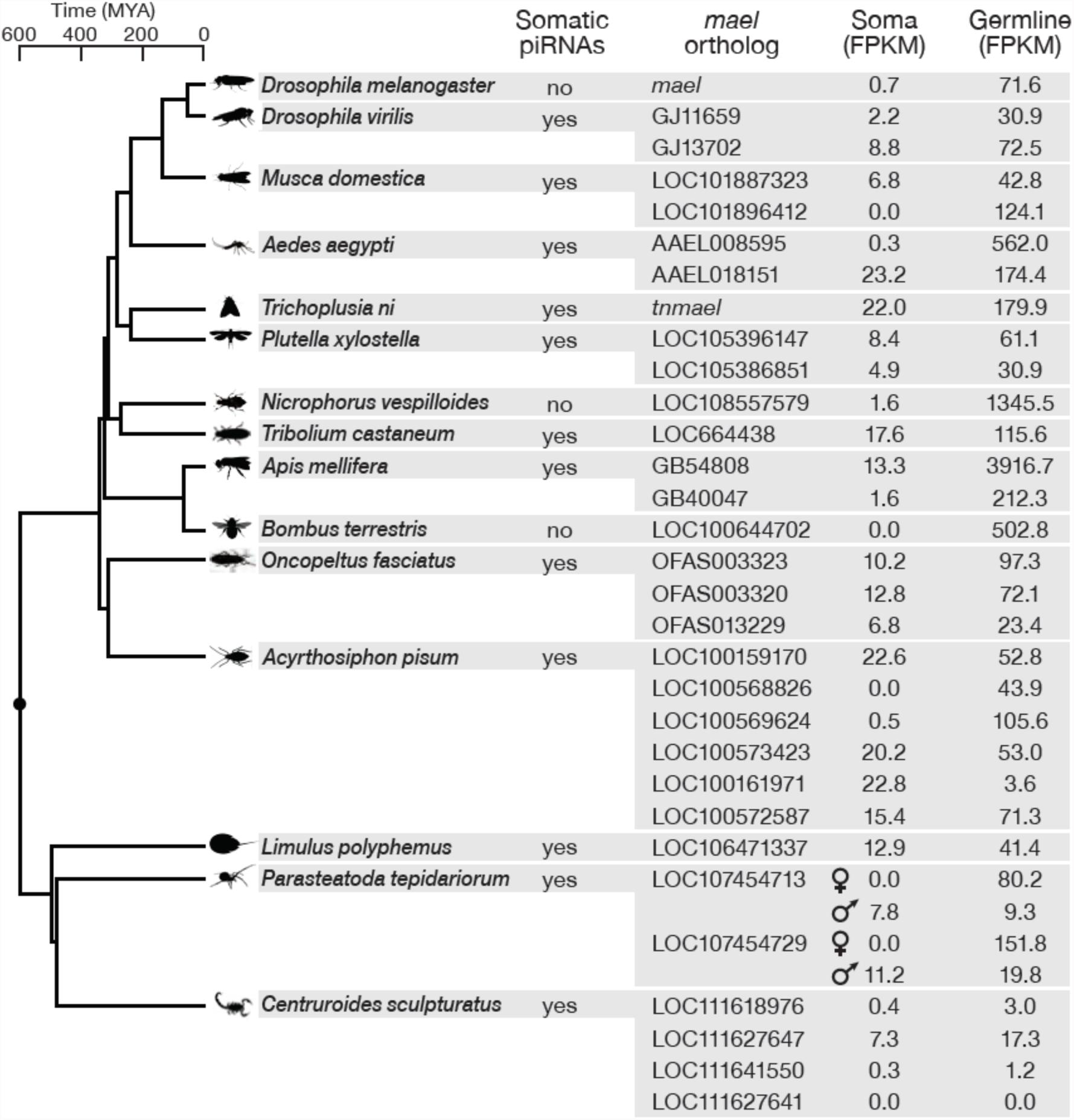
The expression of Mael orthologs in somatic and germline tissues of arthropods correlates with the presence of piRNAs. mRNA expression data are in FPKM from females, except for *Parasteatoda tepidariorum*, where females and males differ in the abundance of somatic piRNAs.

## SUPPLEMENTAL TABLE TITLES AND LEGENDS

**Table S1, related to Figure 6.** Relative White protein function, measured by absorbance at 480 nm of acidic ethanol-extracted eye pigment, in *w^m4^* adult flies. Data are mean ± S.D. (*n* = 3). *p*-values are compared to the *w^m4^* control.

| **white source** | **Mutation** | **Relative OD480** | **p-value** |
| --- | --- | --- | --- |
| $w^{m4}$ | none | $1.0 \pm 0.5$ | control |
| $w^{m4}$ | *mael*<sup>M391/+</sup> | $99 \pm 6$ | $<10^{-4}$ |
| $w^{m4}$ | *mael*<sup>r20/+</sup> | $80 \pm 10$ | $10^{-4}$ |
| $w^{m4}$ | *mael*<sup>M391/r20</sup> | $140 \pm 6$ | $<10^{-4}$ |
| $w^{m4}$ | *Su(var)2-5*<sup>5/+</sup> | $121 \pm 6$ | $<10^{-4}$ |
| $w^{m4}$ | *Su(var)3-9*<sup>2/+</sup> | $148 \pm 3$ | $<10^{-4}$ |

**Table S2, related to Figure S5.**
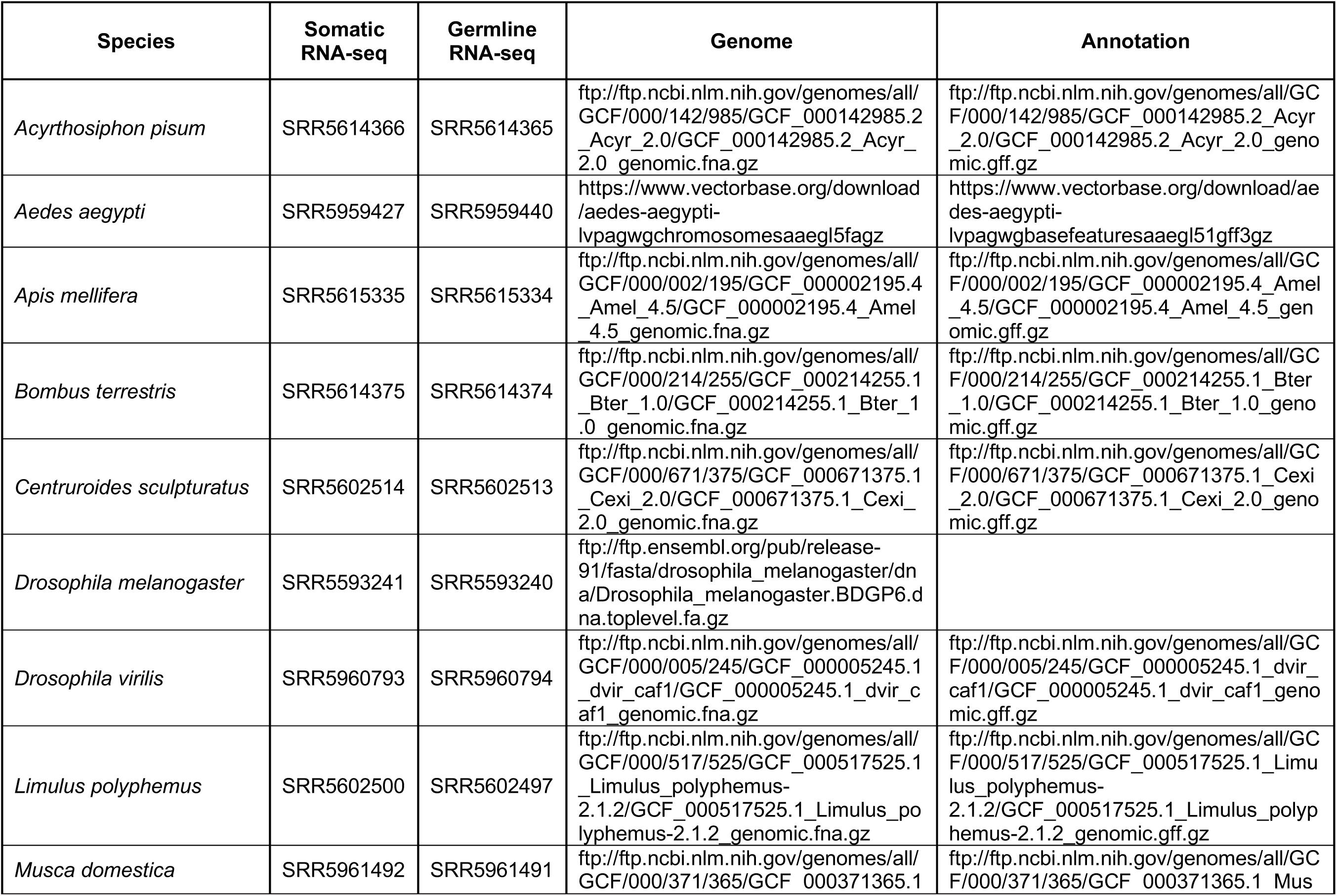
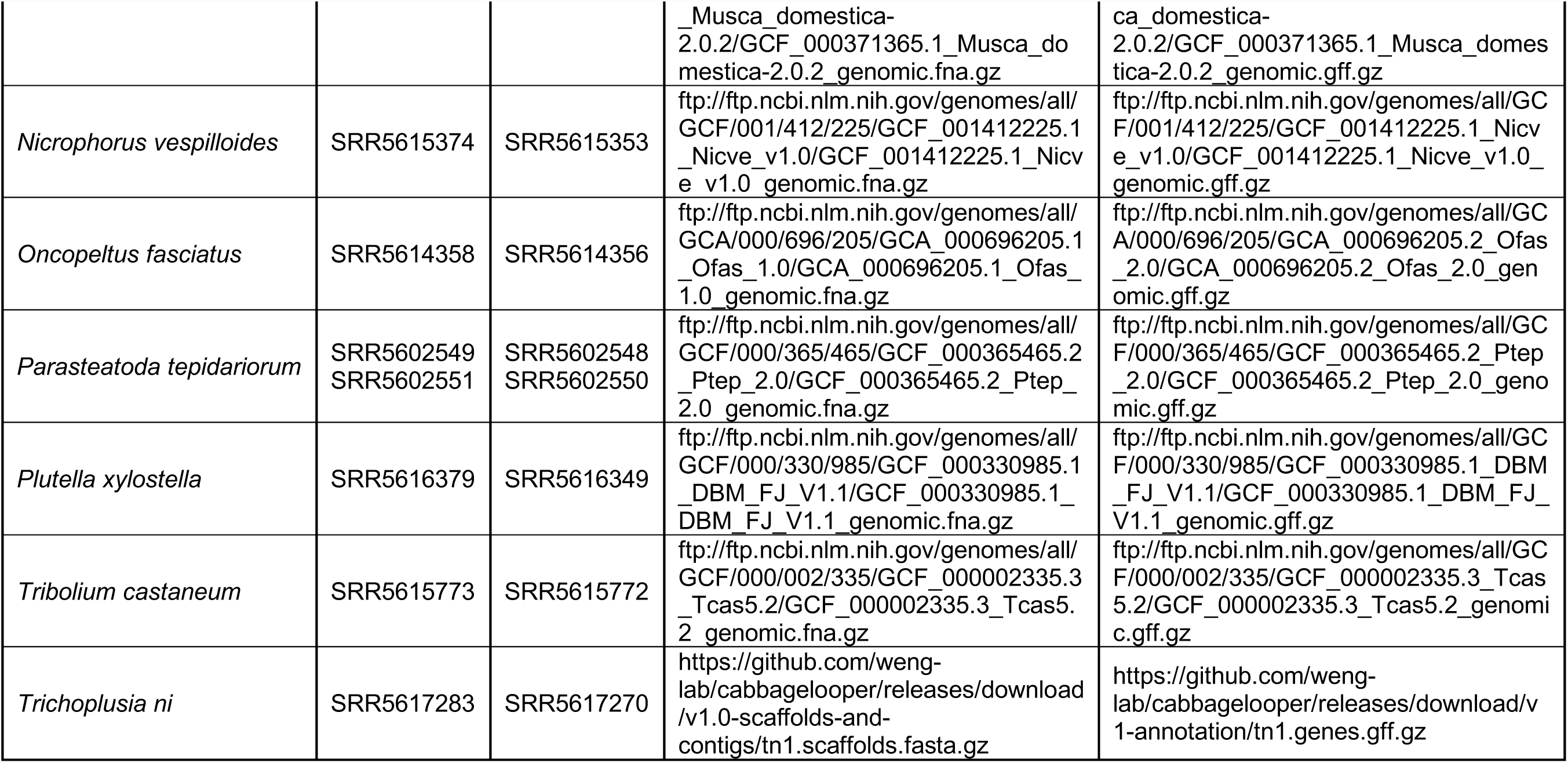
Data sets, genomes, and annotations used to analyze expression of Mael orthologs in 15 arthropods.

## STAR METHODS

### KEY RESOURCES TABLE

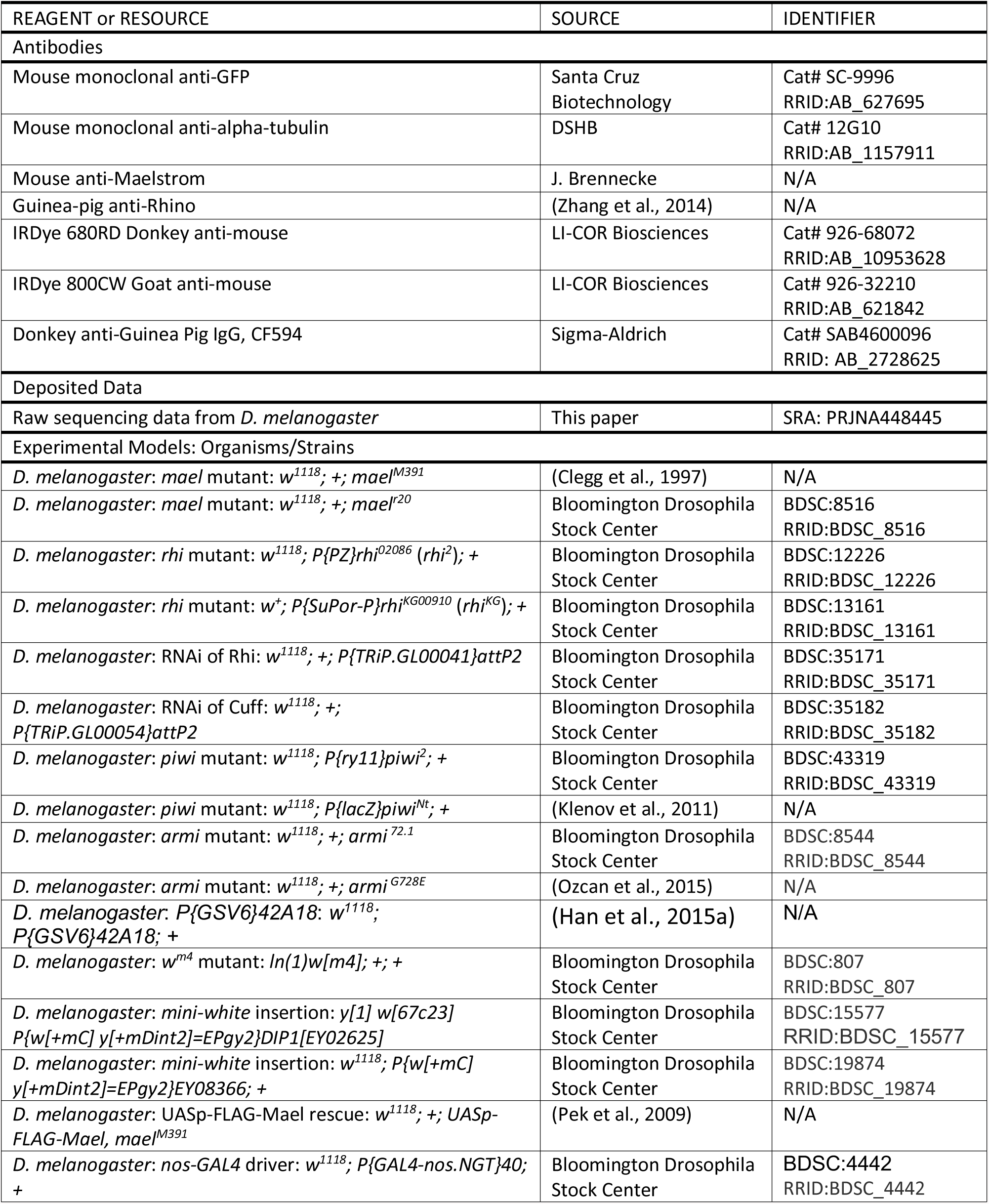
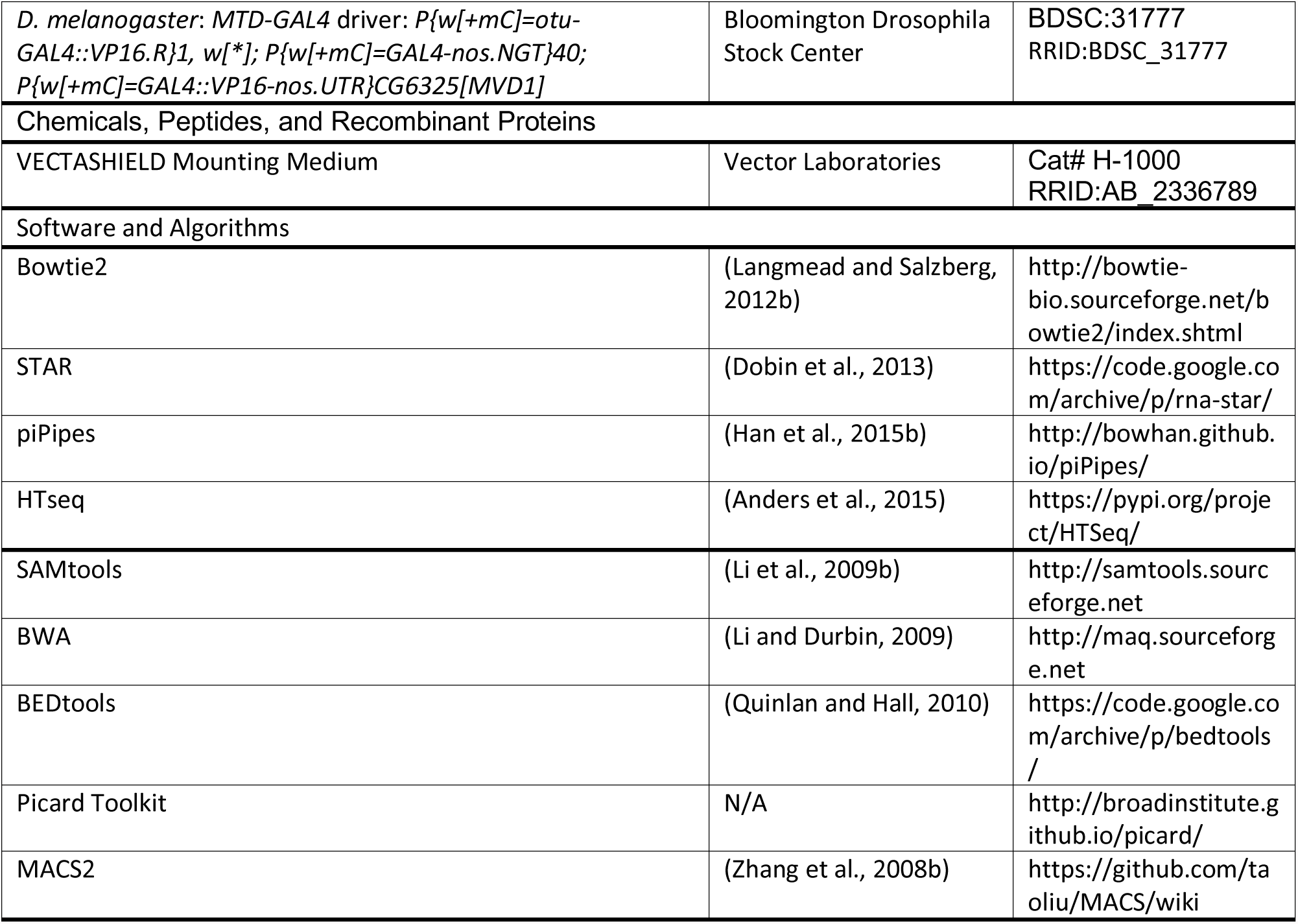

### CONTACT FOR REAGENT AND RESOURCE SHARING

Further information and requests for resources and reagents should be directed to, and will be fulfilled by, the Lead Contact, Phillip D. Zamore.

### METHOD DETAILS

#### *Drosophila* Stocks

Fly stocks were maintained at 25°C. All strains were in the *w^1118^* background except *w^+^*; *rhi^KG^*. Both the *mael^M391^* and *mael^r20^* alleles were backcrossed to *w^1118^* for five generations before use to minimize genetic background effects. The *armi^G728E^* allele was outcrossed for six generations, and the FRT site sequences found in the original line removed. The *P{GSV6}42A18* transgene derives from *P{GSV6}^GS13456^* (Chendrimada et al., 2005) and is located at Chr2R: 6,460,398-6,460,415 (dm6).

#### General Methods

Before dissection, flies were isolated 0–3 days after eclosion and given yeast paste for two days. Fly ovaries were then dissected and collected in 1× phosphate-buffered saline [pH 7.4] (1×PBS) (137 mM NaCl, 2.7 mM KCl, 10 mM Na_2_HPO_4_, 1.8 mM KH_2_PO_4_) cooled on ice. Ovaries were then washed once with ice-cold 1×PBS and then used for subsequent experiments.

#### Western Blotting

Ovary lysate was prepared as described (Li et al., 2009a) with modifications. After 1×PBS was removed, the ovaries were homogenized with a plastic pestle (Fisher Scientific, #12141364) in ice-cold lysis buffer (for each 100 mg ovaries, 100 µl of 100 mM potassium acetate, 30 mM HEPES-KOH [pH 7.4], 2 mM magnesium acetate, 1 mM dithiothreitol (DTT)) containing 1 mM AEBSF (4-(2-aminoethyl)benzenesulfonyl fluoride hydrochloride; EMD Millipore, #101500), 0.3 µM Aprotinin (Bio Basic Inc, #AD0153), 20 µM Bestatin (Sigma Aldrich,#B8385), 10 µM E-64 ((1S,2S)-2-(((S)-1-((4-Guanidinobutyl)amino)-4-methyl-1-oxopentan-2-yl)carbamoyl)cyclopropanecarboxylic acid; VWR, #97063), and 10 µM Leupeptin (Fisher Scientific, #108975). Lysate was centrifuged at 13,000 × *g* for 30 min at 4°C and an equal volume of 2× loading dye (100 mM Tris-HCl [pH 6.8], 4% (w/v) SDS, 0.2% (w/v) bromophenol blue, 20% (v/v) glycerol, and 200 mM DTT) was added to the supernatant and heated to 95°C for 5 min. The lysate was resolved through a 4–20% gradient polyacrylamide/SDS gel electrophoresis (Bio-Rad Laboratories, #5671085). After electrophoresis, proteins were transferred to a 0.45 µm pore polyvinylidene difluoride membrane (Millipore, #IPVH00010), the membrane blocked in Blocking Buffer (Rockland Immunochemicals, #MB-070) at room temperature for 1 h and then incubated overnight at 4°C in 1:1 Blocking Buffer:1 × TBST [pH 7.5] (50 mM Tris-HCl [pH 7.5], 150 mM NaCl, 0.1% Tween 20 (v/v)) containing primary antibody (anti-GFP, Santa Cruz Biotechnology, #SC-9996, 1:2500 dilution; antia-Tubulin, DSHB, #12G10, 1:50,000 dilution, anti-Mael, gift from Julius Brennecke, 1:2500 dilution). The membrane was washed three time, each for 5 min, with 1× TBST [pH 7.5] at 25°C, incubated in Blocking Buffer diluted 1:1 in 1× TBST [pH 7.5] and containing secondary antibody (donkey anti-mouse IRDye 680RD, LICOR Biosciences, #926-68072, 1:10,000 dilution; goat anti-mouse IRDye 800CW, LICOR Biosciences, #926-32210, 1:10,000 dilution) for 1 h at room temperature in the dark, and washed five times for 10 min each with 1× TBST [pH 7.5] at room temperature in the dark. Signal was detected using an Odyssey Infrared Imaging System. Data were obtained from three independent biological replicates. Quantification was performed using Image Studio v4.0.21 (LI-COR). *p*-values were measured using an unpaired, two-tailed t-test.

#### Immunohistochemistry and Microscopy

Immunohistochemistry and microscopy was performed as described (Li et al., 2009a). After ovaries were teased apart using a pipette, they were fixed in 4% formaldehyde in 15 mM piperazine-N,N′-bis(2-ethanesulfonic acid) (PIPES) [pH 7.4], 80 mM KCl, 20 mM NaCl, 2 mM EDTA, 0.5 mM EGTA, and 1 mM DTT (Buffer A) for 10 min, rotating at room temperature. Afterwards, ovaries were washed twice for 15 min each at room temperature in Buffer A with 0.1% (w/v) Triton X-100. The ovaries were washed again twice for 15 min each at room temperature in Buffer A with 10% (v/v) normal donkey serum (Sigma, #D9663) and 0.1% (w/v) Triton X-100. The ovaries were then incubated with anti-Rhi (gift from William Theurkauf Zhang et al., 2014; 1:1000 dilution) in the previous buffer rotating at 4°C overnight.

The next day, the ovaries were washed four times for 30 min each at room temperature in Buffer A with 2 mg/ml BSA and 0.1% (w/v) Triton X-100, followed by a 30 min wash at room temperature in Buffer A with 10% (v/v) normal donkey serum and 0.1% (w/v) Triton X-100. The ovaries were then incubated in the dark, rotating overnight at 4°C with secondary antibody (Donkey anti-Guinea Pig IgG, CF594, Sigma-Aldrich, #SAB460009, 1:100 dilution) in Buffer A with 10% (v/v) normal donkey serum and 0.1% (w/v) Triton X-100.

The next day, ovaries were washed twice for 30 min each at room temperature in in Buffer A with 0.1% (w/v) Triton X-100, followed by a 1 min wash at room temperature with Buffer A. Ovaries were then mounted using VECTASHIELD Mounting Medium (Vector Laboratories, #H-1000) and covered with a 0.13–0.17 mm thick cover slip (VWR #48393 106). Images were captured using a Leica TCS SP5 II Laster Scanning Confocal Microscope using the 63× objective with Olympus Immersion Oil Type-F (Thorlabs #MOIL-30). Images presented in the same figure were acquired at the same settings.

#### Female Fertility Assay

Female fertility was tested essentially as described (Li et al., 2009a). Eight female virgins were mated to four Oregon R virgin males in a small cage with a 60 mm diameter grape juice agar plate dabbed with yeast paste at 25°C. All flies were collected <1-day posteclosion. After two days, the first plate was discarded and replaced with a fresh plate. Plates were then changed and scored every subsequent day. The number of total eggs, eggs per female per day, and the dorsal appendage phenotype of embryos were scored every 24 h and the number of eggs that hatched were scored 48 h after the plate was changed. Fertility was recorded for 12 days and at least two independent biological replicates were conducted for each genotype.

#### Eye Pigment Assay

Ethanol-based pigment extraction and quantification was performed essentially as described (Sun et al., 2004). Briefly, 10 females, 3–5 days post-eclosion, were collected and their heads photographed and dissected. Heads were homogenized in 0.5 ml of 0.01 M HCl in ethanol. The homogenate was incubated at 4°C rotating overnight, warmed to 50°C for 5 min, and centrifuged at 13,000 × *g* for 10 min at 25°C. The supernatant was collected, and absorbance at 480 nm (A480) was measured. Background from *w^1118^* female heads was subtracted from each measurement. *p*-values were measured using an unpaired, two-tailed t-test.

#### Construction and Analysis of sRNA-Seq Libraries

Small RNA libraries were constructed as described (Han et al., 2015a). In summary, total RNA (50 µg) was extracted using mirVana miRNA Isolation Kit (Life Technologies, #AM1561) and purified by 15% urea polyacrylamide gel electrophoresis (PAGE), selecting for 18–30 nt small RNAs. Half of the purified sRNAs were oxidized with NaIO4 was used to deplete miRNAs and enrich for siRNAs and piRNAs (Li et al., 2009a). To reduce ligation bias, a 3′ adaptor with three random nucleotides at its 5′ end was used (5′-rApp NNN TGG AAT TCT CGG GTG CCA AGG /ddC/−3′). 3′ adaptor was ligated using truncated, K227Q mutant T4 RNA Ligase 2 (made in lab) at 25°C for ≥16 h, sRNAs precipitated, and size selected as described in (Li et al., 2009a). To exclude 2S rRNA from sequencing libraries, 10 pmol 2S blocker oligo was added before 5′ adaptor ligation (Wickersheim and Blumenstiel, 2013). 5′ adaptor was added using T4 RNA ligase (Life Technologies, #AM2141) at 25°C for 2.5 h, followed by reverse-transcription using AMV reverse transcriptase (New England Biolabs, #M0277L) and PCR using AccuPrime *Pfx* DNA Polymerase (Invitrogen, #12344-024). Small RNA-seq libraries for three independent biological replicates were sequenced using a NextSeq500 (Illumina) to obtain 75 nt single-end reads.

sRNA-seq analysis was performed with piPipes (v1.4; Han et al., 2015b). Barcodes were sorted allowing one mismatch, and the 3′ adaptors, including the three random nucleotides, were identified and removed using the first ten nucleotides, allowing one mismatch. After adaptor removal, reads containing one or more nucleotides with Phred score <5 were discarded. sRNAs were first aligned to rRNA or miRNA hairpin sequences using Bowtie2 (v2.2.0; Langmead and Salzberg, 2012b). Unaligned reads were mapped to the genome and 23–29 nt RNAs (fly piRNAs) were kept for analyses. The number of piRNAs overlapping each genomic feature (genes, transposons, and piRNA producing loci) were apportioned by the number of times they aligned to the genome.

Oxidized sRNA libraries are enriched with piRNAs. Therefore, to compare piRNA abundances across different oxidized libraries, we calibrated oxidized to unoxidized libraries. Because paired oxidized and unoxidized sRNA libraries were created from the same source, the subset of piRNA species should remain constant between the two libraries. First, unoxidized libraries were normalized to sequencing depth (ppm). Next, we identified all the uniquely mapping piRNA species (piRNAs that shared the exact nucleotide sequences) that were shared between at least two of the three replicates of paired oxidized and unoxidized libraries. Finally, the calibration factor was computed using the ratio between the sums of the normalized abundance in the unoxidized libraries and the abundances in the oxidized libraries,

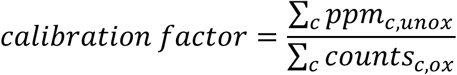

where *c* is the number of common piRNA species between oxidized and unoxidized libraries. piRNAs in the top 10^th^ percentile were excluded to avoid overweighting outliers. The abundance of each piRNA in the oxidized library was calculated by multiplying by the calibration factor.

#### Ping-Pong Analysis

Ping-pong analysis was as described (Zhang et al., 2011). In summary, scores at each 5′-to-5′ distance for two piRNAs were defined as the product of their abundances. The Ping-Pong *Z_10_* score was then the difference of the score at the 5′-to-5′ distance of 10 nt and the mean scores of background distances, divided by the standard deviation of the scores of background distances, defined as distances of 0–9 and 11–20 nt. For analyses including multiply mapping reads, read abundances were apportioned by the number of times the read aligned to the genome.

#### Phasing Analysis

Phasing analysis was as described (Han et al., 2015a). Briefly, sRNA reads were mapped to genome and rRNAs, tRNAs, and snoRNAs were removed. The *Z_x_* score for a distance *x* between the 3′ end of one piRNA to the 5′ end of a downstream piRNA on the same genomic strand was calculated by the difference of the score at the distance *x* and the mean scores of background distances, divided by the standard deviation of the scores of background distances. When *x* = 1, the 5′ end is immediately downstream of the 3′ end (phasing). For analyses including multiply mapping reads, read abundances were apportioned by the number of times the read aligned to the genome. Overlaps at positions 2–20 were used as background to calculate *Z_1_*.

#### Construction and Analysis of RNA-Seq Libraries

RNA-seq libraries were constructed as described (Zhang et al., 2012) with several modifications (Fu et al., 2018a). For ribosomal RNA depletion, RNA was hybridized in 10 µl with a pool of 186 rRNA antisense oligos (0.05 µM/each; Morlan et al., 2012; Adiconis et al., 2013) in 10 mM Tris-HCl [pH 7.4] and 20 mM NaCl and heated to 95°C, then cooled at −0.1°C/sec to 22°C, and finally incubated at 22°C for 5 min. Ten units of RNase H (Lucigen, #H39500) were added and incubated at 45°C for 30 min in 20 µl containing 50 mM Tris-HCl [pH 7.4], 100 mM NaCl, and 20 mM MgCl2. RNA was then treated with 4 units DNase (Thermo Fisher, #AM2238) in 50 µl at 37°C for 20 min. After DNase treatment, RNA was purified using RNA Clean & Concentrator-5 (Zymo Research, #R1016). RNA-seq libraries were sequenced using a NextSeq500 (Illumina) to obtain 75 + 75 nt, paired-end reads.

RNA-seq analysis was performed with piPipes (v1.4; Han et al., 2015b). Barcodes were sorted allowing one mismatch and were identified and removed using the first ten nucleotides, allowing one mismatch. RNAs were first aligned to rRNA sequences using Bowtie2 (v2.2.0; Langmead and Salzberg, 2012b). Unaligned reads were then mapped using STAR to the fly genome (v2.3.1; Dobin et al., 2013). Counts were produced using the “strict” option on HTseq (v0.6.1; Anders et al., 2015).

#### Construction and Analysis of ChIP-Seq Libraries

ChIP-seq libraries were constructed as described (Zhang et al., 2014) with several modifications. Briefly, ∼100 µl ovaries per library were first crosslinked with 2% formaldehyde for 10 min rotating at 25°C in Robb’s medium (100 mM HEPES [pH 7.4], 55 mM sodium acetate, 40 mM potassium acetate, 100 mM sucrose, 10 mM glucose, 1.2 mM MgCl_2_, 1 mM CaCl_2_, 1mM DTT, 1 mM AEBSF, 0.3 µM Aprotinin, 20 µM Bestatin, 10 µM E-64, and 10 µM Leupeptin). Crosslinking was quenched by adding Glycine to a final concentration of 120 mM and for 5 min rotating at 25°C. The ovaries were then washed twice with TBS (50 mM Tris-HCl [pH 7.5], 150 mM NaCl), and twice with ChIP lysis buffer (50 mM HEPES-KOH [pH 7.5], 140 mM NaCl, 1% [v/v] Triton X-100, 0.1% [w/v] Na-Deoxycholate, 0.1% [w/v] SDS).

Ovaries were then sonicated in sonication buffer (1% [w/v] SDS, 10 mM EDTA, 50 mM Tris-HCl [pH 8.0], 1mM DTT, 1 mM AEBSF, 0.3 µM Aprotinin, 20 µM Bestatin, 10 µM E-64, and 10 µM Leupeptin) using an E220 Evolution Focused-ultrasonicator (Covaris) with Duty cycle: 5%, Intensity: 140 watts, Cycles per burst: 200, Temperature: <10°C, Time: 20 min. The sonicated lysate was centrifuged at 13,000 × *g* for 15 min at 4°C. Supernatant was diluted 7-fold with dilution buffer (20 mM Tris-HCl [pH 7.5], 167 mM NaCl, 1.2 mM EDTA, 0.01% [w/v] SDS, 1% [v/v] Triton X-100, 1 mM DTT, 1mM AEBSF, 0.3 µM Aprotinin, 20 µM Bestatin, 10 µM E-64, and 10 µM Leupeptin) and incubated overnight rotating at 4°C with antibody (anti-Rhi or Pre-Immune Serum, gift from William Theurkauf, 20 µl; anti-HP1a, DSHB, #C1A9, 5 µg; normal mouse IgG, Abcam, #ab188776, 5 µg; anti-H3K9me3, Abcam, #ab8898, 10.5 µg; anti-H3K4me3, Abcam, #ab8580, 10.5 µg; anti-Histone H3, Abcam #ab18521, 10.55 µg) bound to 100 µl of Dynabeads Protein A/G (Life Technologies, #10002D/#10004D).

The beads were then washed 2×5 min each with 500 µl of the following buffers: Wash buffer A (20 mM Tris-HCl [pH 8.0], 2 mM EDTA, 0.1% [w/v] SDS, 1%[v/v] Triton X-100, 150 mM NaCl), Wash buffer B (20 mM Tris-HCl [pH 8.0], 2 mM EDTA, 0.1% [w/v] SDS, 1%[v/v] Triton X-100, 500 mM NaCl), Wash buffer C (10 mM Tris-HCl [pH 8.0], 1 mM EDTA, 1% [v/v] NP-40, 1% [w/v] Na-deoxycholate, 0.25 M LiCl) and Wash buffer D (10 mM Tris-HCl [pH 8.0], 1 mM EDTA). All wash buffers also contained 1mM DTT, 1 mM AEBSF, 0.3 µM Aprotinin, 20 µM Bestatin, 10 µM E-64, and 10 µM Leupeptin. Beads were then treated with 20 µg/ml RNase A (Fisher Scientific, #FEREN0531) To reverse crosslink and remove protein, beads were incubated overnight at 65°C with 200 µg/ml Proteinase K (Life Technologies, #25530015) in 2×Proteinase K Buffer (200 mM Tris-HCl [pH 7.5], 2 mM EDTA [pH 8.0], and 1% SDS (w/v). Finally, DNA was purified using phenol:chloroform [pH 8.0] and the library was prepared by sequentially performing end-repair, A-tailing, Y-shaped adaptor ligation, and PCR amplification as described (Zhang et al., 2012).

Barcodes were sorted allowing one mismatch and were identified and removed using the first ten nucleotides, allowing one mismatch. For H3K9me3, HP1a, and Rhi ChIP-seq libraries, reads were mapped on the genome using Bowtie2 (v2.2.0; Langmead and Salzberg, 2012b). Unmapped reads were removed using SAMtools (v0.1.19; Li et al., 2009b; Li, 2011) and a mapping *q*-score of 10 was used to identify uniquely mapping reads. We used a 1 kbp sliding window with a 500 bp step over the genome to compute a signal for each chromosome arm. For each chromosome arm, the counts for each bin were normalized using the total number of reads mapping to the chromosome. Unique reads were also mapped to genomic features (genes, transposons, and piRNA producing loci) using STAR (v2.3.1; Dobin et al., 2013) and counts were produced using the “strict” option on HTseq (v0.6.1; Anders et al., 2015). Reads were normalized to sequencing depth.

H3K4me3 ChIP-seq libraries were analyzed using the psychENCODE pipeline (PsychENCODE et al., 2015). Briefly, reads were mapped to the genome using BWA (Li and Durbin, 2009). SAMtools (v0.1.19; Li et al., 2009b; Li, 2011) and Picard Toolkit (Ioannidis, 2005)Ioannidis, 2005 were used to remove improperly paired reads and PCR duplicates. MACS2 (Zhang et al., 2008b) was used to called peaks. We used BEDtools (v2.26.0; Quinlan and Hall, 2010) to merge peaks from all replicates for each genotype to create a consensus set of peaks. The number of reads overlapping each peak was computed using BEDtools and reads were normalized to sequencing depth.

#### Construction and Analysis of GRO-Seq Libraries

GRO-seq libraries were constructed as described (Wang et al., 2015), and analyzed with piPipes (v1.4; Han et al., 2015b). Briefly, 0–2-day-old female flies were given yeast for 2 days before their ovaries were dissected. One hundred pairs of ovaries were homogenized in 350 µl HB35 buffer (15 mM HEPES KOH [pH 7.5], 10 mM KCl, 2.5 mM MgCl2, 0.1 mM EDTA, 0.5 mM EGTA, 0.05% [v/v] NP 40, 0.35 M sucrose, 1 mM DTT, 1 mM AEBSF, 0.3 µM Aprotinin, 20 µM Bestatin, 10 µM E-64, and 10 µM Leupeptin) with a Dounce homogenizer using pestle B (Sigma Aldrich, #D8938). Nuclei were purified by passing twice through sucrose cushions that contain 800 µL HB80 buffer (15 mM HEPES KOH [pH 7.5], 10 mM KCl, 2.5 mM MgCl2, 0.1 mM EDTA, 0.5 mM EGTA, 0.05% [v/v] NP 40, 0.80 M sucrose, 1 mM DTT, 1 mM AEBSF, 0.3 µM Aprotinin, 20 µM Bestatin, 10 µM E-64, and 10 µM Leupeptin) on the bottom phase and 350 µl HB35 buffer on the top. Nuclei were washed once with 500 µl freezing buffer (50 mM Tris-HCl, pH 8.0, 40% [v/v] glycerol, 5 mM MgCl2, 0.1 mM EDTA, 1 mM dithiothreitol, 1 mM AEBSF, 0.3 µM Aprotinin, 20 µM Bestatin, 10 µM E-64, and 10 µM Leupeptin) and frozen in liquid nitrogen with 100 µL freezing buffer. To carry out nuclear run on assay, 100 µL freshly prepared reaction buffer (10 mM Tris-HCl, pH 8.0, 5 mM MgCl2, 300 mM KCl, 1% [w/v] sarkosyl, 500 µM ATP, 500 µM GTP, 500 µM Br UTP, 2.3 µM CTP, 1 mM dithiothreitol, 20 U RNasin Plus RNase Inhibitor (Promega, #N2615) was added to nuclei and incubated at 30°C for 5 min. RNA was extracted using Trizol (Invitrogen, #15596). Nascent RNAs with Br UTP incorporated were enriched by immunoprecipitation using anti 5 bromo 2′ deoxyuridine antibody (Fisher Scientific, #50175223) as described (Shpiz and Kalmykova, 2014), followed by rRNA depletion using RNase H, fragmentation, and library construction as in RNA-seq library preparation (Zhang et al., 2012).

#### Analysis of Mael Orthologs in Arthropods

Mael orthologs in arthropods were identified using either available genome annotation (Table S2; *Musca domestica, Plutella xylostella, Nicrophorus vespilloides, Limus polyphemus, Parasteatoda tepidariorum, Centruroides sculpturatus, Trichoplusia ni*) or OrthoDB v9.1 (group EOG091G0BYM; Zdobnov et al., 2017; *Drosophila virilis, Aedes aegypti, Tribolium castaneum, Apis mellifera, Bombus terrestris, Oncopeltus fasciatus, Acyrthosiphon pisum*). RNA-seq data sets (Table S2; Lewis et al., 2018; Fu et al., 2018b) were first aligned to the corresponding organism’s ribosomal RNA sequences (SILVA database; Quast et al., 2013) using Bowtie2 (v2.2.0; Langmead and Salzberg, 2012a). Unaligned reads were then mapped to the corresponding organism’s genome assembly (Table S2) using STAR (v2.3.1; Dobin et al., 2013). Sequencing depth and gene quantification was calculated with Cufflinks (v2.1.1; Trapnell et al., 2010).

#### Data Availability

Sequencing data are available from the NCBI BioProject Archive using accession number PRJNA448445.

